# Zinc-starved Brassicaceae Plants Secrete Peptides that Induce Root Expansion

**DOI:** 10.1101/2024.06.11.598559

**Authors:** Sarah P. Niehs, Jakub Rajniak, Anna Johnson, Diego L. Wengier, Elizabeth S. Sattely

## Abstract

Zinc (Zn) deficiency is recognized as a global crisis as it is observed in half of all agricultural soils. However, the molecular mechanisms that drive plant physiological responses to soil Zn deficiency are not well understood. We used an untargeted metabolomics approach to search for metabolites exuded from roots during Zn deficiency stress, which led to the discovery of a collection of secreted small defensin-like peptides in *Arabidopsis thaliana* (named Zinc-Deficiency Responsive Peptides (ZDRPs)). Phylogenetic analysis and untargeted metabolomics revealed ZDRPs in at least eleven accessions of *A. thaliana* and nine members of the Brassicaceae family. Analysis of *Arabidopsis* gene mutants and overexpressing lines, in combination with chemical complementation experiments, unveiled a critical role of these peptides in plant root growth. We hypothesize that Brassicaceae secreted peptides enable plants to expand their root mass to reach Zn-rich soil layers and optimize Zn uptake. These data reveal a critical relationship between plant survival, Zn status, root morphology and peptide production. Taken together, our results expand our knowledge regarding micronutrient deficiency responses in plants and could enable in engineering approaches to make plants more resilient to low Zn conditions.

**Significance:** Zinc deficiency is the most abundant micronutrient deficiency affecting about 50% of arable lands thus presenting a high burden for plant health and agriculture globally. In this study, we reveal a metabolic strategy by Brassicaceae to deal with low Zn concentrations. We characterize the role of peptides expressed upon zinc deficiency in a variety of important crop plants. The discovery of a cryptic class of peptides that are made by plant roots specifically suffering from Zn deficiency provides critical insight into the molecular mechanisms by which plants dynamically acclimate to nutrient-limited soils. The identification of peptides actively secreted by zinc-deprived plants has translational value for sustainable agriculture, human health, and bioengineering approaches to enable tolerance to low zinc.

## Introduction

Plants face nutrient deficiencies during their life cycle and differ in their ability to cope with non-ideal growth conditions. While much is known about how plants respond to macronutrient deficiencies, micronutrient deficiency responses are less well understood (1, 2).

Fifty percent of arable land shows signs of zinc (Zn) deficiency making it the most ubiquitous micronutrient deficiency problem in the world (3). Zn deficiency can occur naturally in calcareous soils, but the application of larger amount of phosphate-containing fertilizers for farming has led to a steep increase in generating additional Zn-deficient soils (3, 4). While many soil macro- and micronutrient deficiencies in plants are commonly addressed by the application of fertilizers, a complementary strategy would be to leverage natural mechanisms evolved in the plant kingdom through plant engineering.

In plants, Zn is essential for numerous core cellular processes (5). Plants suffering from severe Zn deprivation show stunted growth, chlorosis of leaves, sterility, and decreased defense capabilities leading to major losses in agriculture (3). Nonetheless, plants can acclimate to Zn deprivation using various strategies, such as the expression of genes that regulate soil Zn uptake and translocation, strict regulation of Zn-containing enzymes, and release of phytochelators (6). In addition, plant root morphology changes under Zn deprivation have been observed in various plant species (7–9), including extended branching of the root system of *Arabdopsis thaliana* (10).

While some aspects of Zn acquisition such as Zn transporters in plants are understood, the molecular mechanisms behind the observed changes in root morphology that drive acclimation to local Zn deficiency and the diversity of strategies that have evolved across the plant kingdom to cope with low Zn availability are not well understood. A critical advance has been the discovery of the F-group of the BASIC LEUCINE-ZIPPER transcription factors bZIP19 and bZIP23 that drive recognition and regulation of molecular plant responses in Zn-deficient soils or growth medium (11). These proteins harbor a Zn-sensing motif compromised of cysteines and histidines that were shown to directly bind two Zn ions per protein (12). Under Zn deprivation, bZIP19 and bZIP23 dimerize and bind to the Zinc Deficiency Response Element (ZDRE) in the promoter sequence of over 83 predicted target genes in Zn-deprived *Arabidopsis thaliana* (13), such as Zn transporter genes, NICOTIANAMINE SYNTHASE genes and cryptic DEFENSIN-LIKE (*DEFL*) peptide genes (14). While some of these target genes have known functions, the roles of many others are still unknown. One of these DEFL peptides have been previously studied in the context of antinematodal activities (15). Another study focused on two *DEFL* peptide genes whose gene expression was upregulated under Zn deficiency (16). An effect on root meristem length and cell number in DEFL mutant lines has also been reported (16).

Since the Zn stores necessary for plant growth reside in the soil, we predicted that small molecules secreted from plant roots could help plants orchestrate responses to Zn deficiency. Interactions of plants with the soil environment can be crucially influenced by plant root exudates. For example, plant root exudates can contain primary metabolites to mobilize metals, secondary metabolites to alter microbial microbial composition (17) and proteins for defense or for nutrient scavenging (18) and are therefore a rich source to study interactions of plants and soil environment. One widespread strategy to deal with many types of metal deficiency and toxicity involves secretion of metal-chelating compounds like nicotianamine (19, 20). These metabolites are not specific to Zn, often having higher affinity to iron (21). This promiscuity in metal recognition can lead to toxic accumulation of high abundant metals from the environment in the host plant (22). In a related investigation of micronutrient deficiency, we used an untargeted metabolomics approach with *Arabidopsis* seedlings to uncover redox active coumarins secreted from plant roots during iron deficiency (23). We reasoned that a similar approach would give valuable insights into the first line metabolic response that are specific to Zn-deficiency stresses, and therefore investigated *A. thaliana* and related plants’ exudates specifically under Zn starvation.

Here, we report the discovery of a collection of small peptides derived from *DEFL* genes that are produced and secreted into the rhizophere in Brassicaceae plants during Zn deficiency. Through the combination of phenotypic analysis of gene knockout mutants and overexpression lines, along with chemical complementation studies and metabolic profiling, we show that these <u>Z</u>inc-<u>D</u>eficiency <u>R</u>esponsive <u>P</u>eptides (ZDRPs) function in root growth. Taken together, we propose ZDRPs enable root foraging for nutrition when soil Zn reservoirs are low.

## Results

### Distinct defensin-like peptides are produced under Zn deficiency and actively secreted into soil

To uncover the metabolic response of plants under Zn deficiency, we first established a system that would enable root exudate sampling, medium exchange, and easy separation of aerial from root tissue in a sterile environment (Figure 1A). Since agar and other solid media contain sufficient amounts of Zn for Brassicaceae to survive (24), we employed a liquid hydroponics set-up with the model plant *Arabidopsis thaliana* ecotype Columbia-0 (Col-0) (SI methods). Sterile stratified plant seeds were placed on PTFE mesh floating over Murashige-Skoog nutrient solution containing no zinc (-Zn) or sufficient amounts of Zn (1.94 mg/L, basal). WT seedlings grown under limiting Zn for 28 days had a higher root dry weight (DW) than plants from basal medium, but lower shoot DW (Figure 1B). Under Zn deprivation, increased total root length has been observed for barley (7) as well as higher primary root length for rice (8) and sorghum (9). In addition, mild Zn deficiency has been shown to lead to a more highly branched root system in *A. thaliana* (10). We observed a yellowing of leaves of plants grown under -Zn in the hydroponics set-up, a sign of plant Zn deficiency in the field (Figure 1B). These data indicate our hydroponics seedling growth assay – with tight control of Zn content – can induce many of the Zn deficiency symptoms observed in plants grown in natural soils.

**Figure 1.**
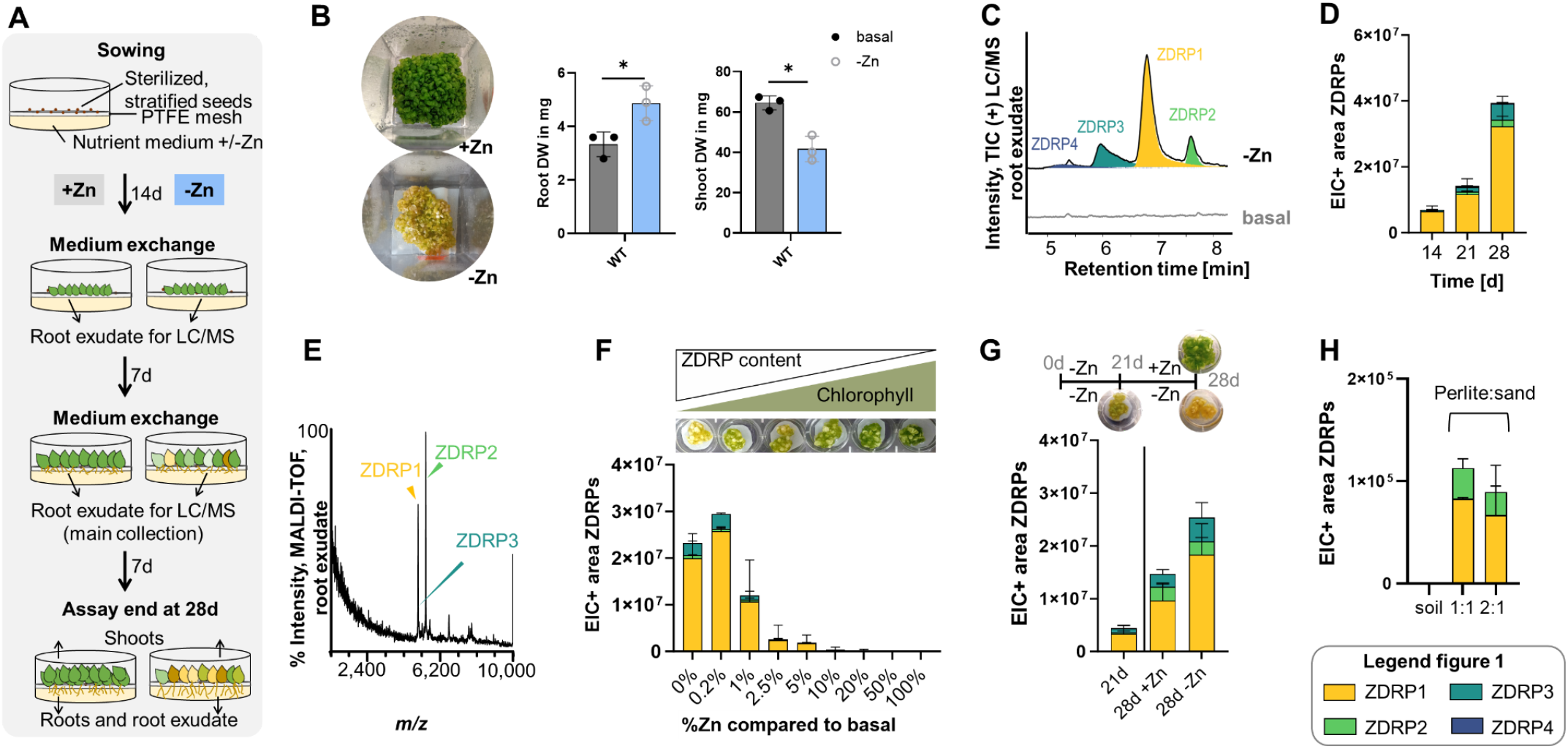
Identification of the metabolic plant response to Zn deficiency. A) Hydroponics assay set-up to determine the metabolic plant response under Zn deficiency. Each datapoint represents one well. One well has 4 mg of seed weight at the start of the experiments (ca. 250 seeds). B) Chlorosis phenotype observed in -Zn plants as well as root and shoot dry weights of 28-day old seedlings grown under -Zn or basal conditions. P-values indicated. Statistical analysis, unpaired t-test. C) Metabolic profiles of root exudates from *A. thaliana* Col-0 grown under Zn-deficient conditions (-Zn) or basal conditions detected using LC/ESI-MS after 28 days. Extracted ion chromatograms of ZDRP1-4 are highlighted. D) LC/MS analysis of root exudates from plants grown under -Zn conditions over time. E) MALDI-TOF-MS analysis of root exudate harvested from *A. thaliana* Col-0 after Zn deprivation. F) LC/MS analysis of root exudates from of *A. thaliana* plants grown under various Zn concentrations after 21 days and chlorosis phenotype under various Zn concentrations after 28 days (0-100% of basal concentration). G) LC/MS analysis of root exudates from *A. thaliana* plants after continuous Zn deprivation for 21 days. After 21 days of growth in media with no added Zn, seedlings were transferred to either Zn deficient or Zn-replete media and ZDRP production was measured at 28 days. H) LC/MS analysis of water extracts from perlite:sand growth medium after 6 weeks of *Arabidopsis* growth. EICs for all observed ZDRPs shown. Individual data points represent plants and root exudates in one well. Error bars represent standard deviations. Replicates per bar: n=3 (1D), n=5-7 (1F), n=3-4 (1G), n=2 (1H). *P*-values: *, ⪬0.05.

To search for a wide range of root metabolites, we collected the plant root exudates at three different time points and measured the crude extracts by reversed phase and HILIC LC/ESI-MS in positive mode (Figure S1A-B). Interestingly, the metabolic profile of -Zn root exudate revealed four distinct mass signatures absent from the +Zn control (reversed phase-LC/MS, Figure 1C, SI methods). These mass signatures were predominantly present in root exudates and only found in trace levels in root and shoot tissue extracts of Zn-deprived plants (Figure S1C). Closer investigations revealed a time-dependent secretion of these compounds from the roots of *A. thaliana* (Figure 1D). We found one mass signature (“ZDRP4”) that accumulates in root exudates only after continuous Zn deprivation for >30 days. Isotopic patterns of all four mass signatures showed multiple charged species (*z* = +5) which would correspond to molecules around 5,300-5,700 Da, common for peptides (Figure S1D). MS/MS of ZDRP1 and ZDRP2 revealed the presence of amino acid fragments corroborating their peptidic nature (Figure S1E). Multiple charged, observed monoisotopic masses were *m/z* 1060.8601 (calculated full-length peptide 5,299.3 Da; ZDRP1), 1135.4746 (calculated 5,672.3 Da, ZDRP2), 1068.2497 (calculated 5,336.2 Da, ZDRP3), 1119.8699 (calculated 5,594.4 Da, ZDRP4). Because of their peptidic nature and their responsiveness to the Zn status of the plant, we named these compounds Zinc-Deficiency Responsive Peptides (ZDRPs).

Next, we employed Matrix-assisted laser desorption ionization time-of-flight mass spectrometry (MALDI-TOF MS) to gain further insight into the full extent of the peptide response (SI methods). MALDI-TOF MS represents a reliable method to detect proteins and peptides as it is a softer ionization method than ESI-MS with minimal instrumental set-up and a wide mass range. We solely observed masses corresponding to ZDRP1, ZDRP2, and ZDRP3 in the -Zn whole root exudate in a mass range from 500 to 10,000 Da (Figure 1E). Taken together, these results reveal that ZDRPs are the dominant metabolite species secreted by *A. thaliana* specifically in response to Zn deprivation.

To study in more detail the dependence of Zn concentrations on peptide production, we employed a Zn gradient ranging from 0% to 100% of the overall basal Zn concentration in MS medium (1.95 mg Zn/L). After 28 days the seedling root exudates were collected and analyzed by LC/ESI-MS. As an indicator for overall plant health, we monitored chlorophyll content by a spectrophotometer assay (Figure S1F, SI methods). Plants grown with 100% Zn were healthy as indicated by their high chlorophyll content with no detectable ZDRP levels (Figure 1F). The peptide levels gradually increased with lowering Zn concentration while the chlorophyll levels decreased. Thus, we found a correlation between ZDRP levels, Zn concentration, and plant health.

To determine whether ZDRP production dynamically changes in response to plant Zn status, we Zn-starved plants for 21 days and then either added fresh media with basal levels of Zn or replenished seedlings with media without Zn. The collected root exudates showed that plants experiencing continuous Zn deprivation gradually increased in ZDRP levels until day 28 while ZDRP levels subsided in the exudates of plants that were replenished with Zn-containing media (Figure 1G and S2A-D). These data suggest that ZDRP production is strictly controlled in *A. thaliana* in response to Zn levels in the growth media.

To test if the observed peptide response is not an artifact of plants grown in hydroponic media, we analyzed exudates from *A. thaliana* cultivated on a solid substrate that mimics soil environments with low Zn. Seedlings were grown for six weeks on high pH soil (where Zn availability is lower) or in nutrient-deficient substrates (sand-soil mixtures, or sand-Perlite ratios). When plant leaves began to show signs of chlorosis, an indicator of micronutrient starvation including Zn, substrate from the rhizosphere was harvested, washed, and the concentrate was subjected to LC/ESI-MS analysis. Only the sand-Perlite substrate yielded detectable levels of ZDRPs (Figure 1H and Figure S2E). ZDRP1 was still the most abundant peptide in line with the hydroponics results, followed by ZDRP2. ZDRP4 was not detectable in artificial soil.

### Untargeted metabolomics unveils link between biosynthetic genes and peptide production

Next, we sought to understand the genetic basis for ZDRP biosynthesis. To begin, we focused on the fragmentation pattern of ZDRP1 obtained from tandem mass spectrometry analysis (MS/MS). We substracted adjacent y-ions values to obtain monoisotopic mass values corresponding to amino acid fragments. Due to ambiguity in the exact mass differences, we determined a partially degenerate sequence for ZDRP1 (Figure 2A and Figure S3). These spectra contained sufficient information to identify the N→C motif F-[VTC]-G-S-[LIND]-A-[LIND] for ZDRP1 with the degenerate sequence indicated in brackets. Applying ScanProsite (25) we searched the *A. thaliana* proteome with the sequence tag which yielded two hits: At3g59930 and At5g33355 for ZDRP1. These two genes belong to Cysteine-Rich Peptide family 0770 (CRP0770) and are predicted to code for DEFL peptides. The CRP0770 family in *A. thaliana* was predicted to be comprised of seven members (Figure 2B) (26).

**Figure 2.**
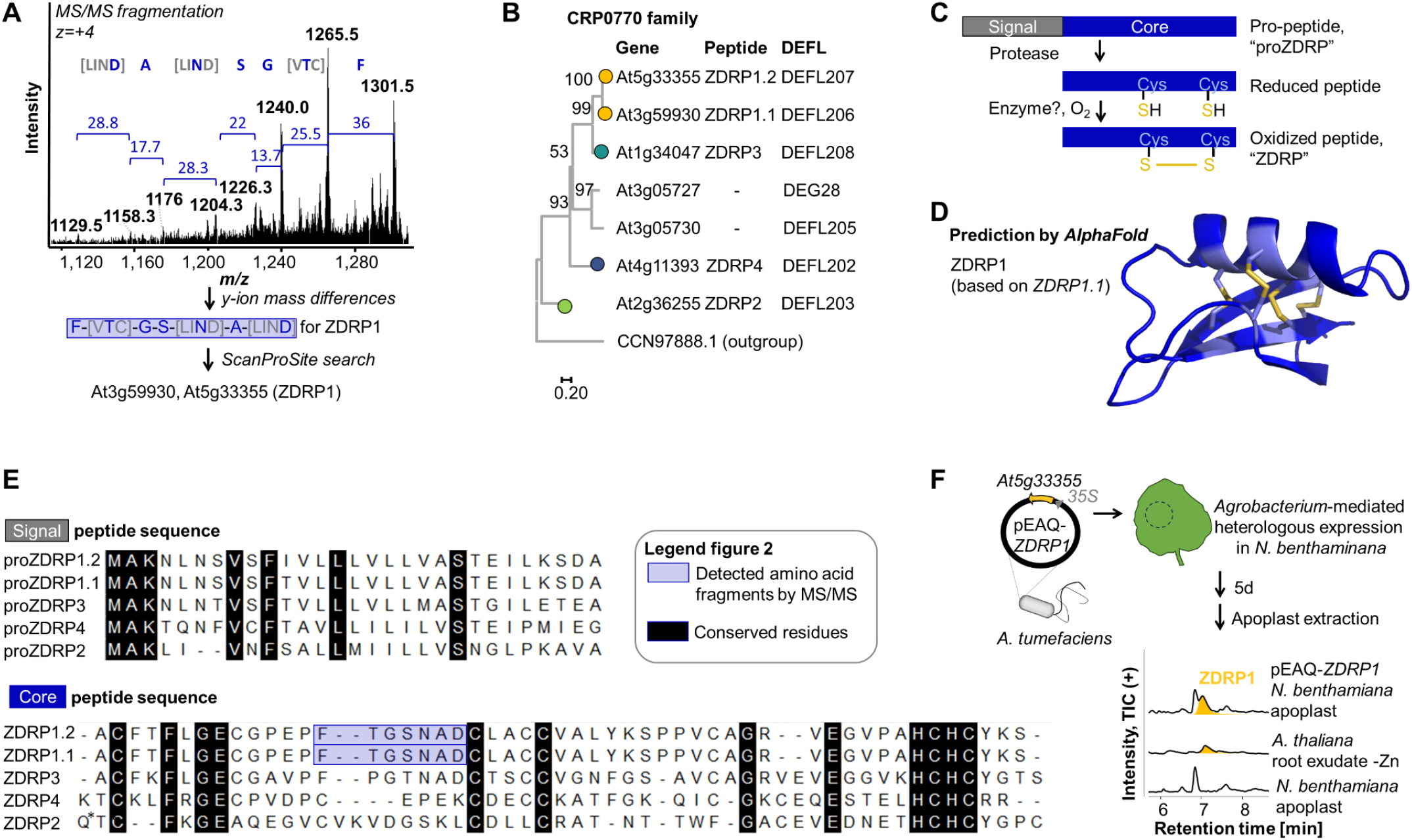
Characterization of ZDRP structure and biosynthesis. A) MS/MS fragmentation of ZDRP1 reveals amino acid fragments that enables identification of ZDRP candidate genes in *A. thaliana* genome. B) Phylogenetic tree of CRP0770 DEFL peptide family constructed with the propeptide sequences. Bootstrap 100. C) Biosynthesis scheme for cysteine-rich small peptides of DEFL class. D) Structure prediction of ZDRP1 core peptide generated by AlphaFold. Core peptide sequence (blue) with disulfide bridges (yellow). E) ZDRP sequences with conserved sites (highlighted). N-terminal Q* cyclized to pyroglutamic acid. Blue, detected amino acid fragments. Abbreviation: pro, propeptide. F) LC/MS analysis of *N. benthamiana* apoplast extracts after *Agrobacterium*-mediated heterologous expression of *ZDRP1*.*2* and root exudate of *A. thaliana* grown under Zn deficiency. EIC of ZDRP1 *m/z* is highlighted. Experiment performed in triplicate.

In addition to MS/MS fragmentation analysis, we examined seven transcriptomics and proteomics datasets dealing with various aspects of Zn deficiency in *A. thaliana* (27–33). All datasets revealed between one to five of the seven predicted CRP0770 *DEFL* peptide genes are induced upon Zn deprivation: At3g59930 (*ZDRP1*.*1*), At5g33355 (*ZDRP1*.*2*), At2g36255 (*ZDRP2*), At1g34047 (*ZDRP3*), and At4g11393 (*ZDRP4*) (Figure S4A). In contrast, the other two members of CRP0770, At3g05727 and At3g05730, are not responsive to Zn deficiency. At3g05727 and At3g05730 share little primary sequence identity with the other five *DEFL* peptide genes (27% by standard Smith-Waterman alignment (34)). Based on their amino acid sequence, At3g05727 and At3g05730 are expected to form three disulfide bridges compared to four to five disulfide bridges in ZDRPs. Their expression pattern in plant tissues does not match that of the other five related *DEFL* peptide genes (26). In light of these stark differences, we hypothesize that At3g05727 and At3g05730 serve other functions unrelated to Zn-deficiency.

The ZDRP family comprises five members with Zn-related functions. *ZDRP1*.*1* and *ZDRP1*.*2* code for the same core peptide sequence (ZDRP1) and only differ in one amino acid near the N-terminus. Their evolution putatively follows a duplication event in the *A. thaliana* genome. ZDRP3 and ZDRP1 are highly related only differing in one amino acid in the signal sequence (Figure 2B). It should be noted that two gene models have been assigned to *ZDRP3*, but our data supports gene model At1g34047.1. In the case of ZDRP2 (At2g36255), we noticed a 17 Da shift between the predicted (5689.4 Da) and observed mass (5672.4 Da). This difference can be attributed to a loss of ammonia following cyclization of the N-terminal glutamine to pyroglutamic acid through spontanous pyroglutamination (35) or enzymatical catalysis by glutaminyl cyclases (36).

We detected *m*/*z* values that match the oxidized form of the peptides with all 8-10 cysteines forming disulfide bonds (Figure 2C). The disulfide bonds could lead to a coiled structure of these DEFL peptides making them highly resistant to breakdown (Figure 2D). Using AlphaFold (37, 38), we noted varying cysteine disulfide patterns in the predicted peptide structures of ZDRPs (Figure S4B-C). All five identified *DEFL* peptide genes code for cysteine-rich (8-10 cysteines) peptides with a secretion signal sequence of 27-29 amino acids and a core sequence of 49-51 amino acids (Figure 2E). There are 12 conserved residues in the core sequences, five of which are cysteines. Interestingly, *ZDRP* genes share a conserved HCHC motif near their C terminus. Besides the HCHC motif, ZDRP sequences are heterogeneous including the number and relative positions of cysteines.

To definitively link these candidate genes to the produced peptides, we used an *Agrobacterium*-mediated transient expression system in *Nicotiana benthamiana* leaves to produce heterologous gene products. We detected the mass signatures in the apoplast of infiltrated leaves that match the -Zn root exudates of *A. thaliana* (Figure 2F and S5). Thus, five members of the *DEFL* CRP0770 gene family in *A. thaliana* can be linked to the four peptides secreted by seedlings under Zn deficiency and no *Arabidopsis*-specific processing or post-translational modification appears to be required to obtain the mature peptide. Transient overexpression in *N. benthamiana* also enabled us to obtain high amounts of peptide used to elucidate the biological function of ZDRPs in experiments described below.

Given the responsiveness between Zn and ZDRPs, we wanted to understand if ZDRPs are part of the known Zn regulatory network. Plant response to Zn deficiency has been identified as a strictly controlled process involving transcription factors bZIP19 and bZIP23 (14). In the original study bZIP19/23 dimerize in the absence of Zn, bind to a 10 bp-long, near-palindromic promoter sequence (Zinc Deficiency Response Element, ZDRE) in the promoter sequence and activate targeted gene expression (14). We analyzed the 3,000 bp region upstream of each gene in the CRP0770 subfamily from *A. thaliana* using a pattern-matching algorithm (39) and found at least one copy of the ZDRE with at most one mismatch from the consensus for all five genes (Figure S6). At3g05727 and At3g05730, the proposed additional members of CRP0770, did not show ZDRE motifs in their upstream region. All in all, ZDRP biosynthetic genes have the landmarks of being targets for upregulation by bZIP19/23 in response to Zn deficiency. Further experiments confirmed this hypothesis (experiments below).

### Conservation and diversification of peptides in response to low Zn in natural variants of *Arabidopsis thaliana* and additional Brassicaceae

As a first step towards examining the ecological role of ZDRPs, we examined their production across *Arabidopsis* ecotypes collected from different locations (North America, Europe, and Asia, Figure S7). Silent or loss-of-function single nucleotide polymorphisms (SNPs) in different ecotypes of *A. thaliana* can reveal genes that are important for survival or provide adaptive advantages in a given habitat. To determine the conservation of Zn-related genes, we investigated the frequency of SNPs in known Zn-related genes. We concentrated on high impact SNPs (loss of start codons, frame shifts, or stop codon gained) rendering the encoded protein non-functional. Applying the POLYMORPH1001 tool (40), we found that there are 36 out of 1,135 ecotypes showing high impact SNPs in *ZDRP* genes. While most mutations occur in *ZDRP3*, we found that all genes accumulate mutations except *ZDRP1*.*1* and transcription factor *bZIP19* which strongly suggests a crucial role in plant survival (Figure S7A). In addition, the duplication of ZDRP1-producing genes and the absence of mutations in *ZDRP1*.*1* indicates a primary role of this specific peptide gene while *ZDRP2-4* might serve secondary or complimentary functions.

To clarify if *ZDRP* mutations can be tied to lower Zn assimilation efficiency of ecotypes, we selected eleven *A. thaliana* ecotypes to screen in a hydroponics set-up for 28 days (Figure 3A). These lines included five with mutations in *ZDRP* genes, one with Zn transporter *IRT3* mutation, one with *bZIP23* mutation, and six lines that have been previously investigated for their response to Zn deficiency (41). Ecotypes showed a wide range of phenotypes under -Zn. Shoots of strongly affected plants under -Zn turned yellow, a sign for severe Zn deficiency (Figure 3A). Ecotype Hau-0 was most affected in the experimental set-up while Kardz-1, Co-1, WI-0, and TÅD plants grown under -Zn were indistinguishable from ones grown under basal conditions. Based on this initial phenotyic assay, we determined that *ZDRP* loss-of-function mutations do not directly lead to significant survival decrease under Zn deficiency. There might also be complementary effects of Zn deficiency-related genes that compensate for the impact of a given one SNP on a Zn-response phenotype.

**Figure 3.**
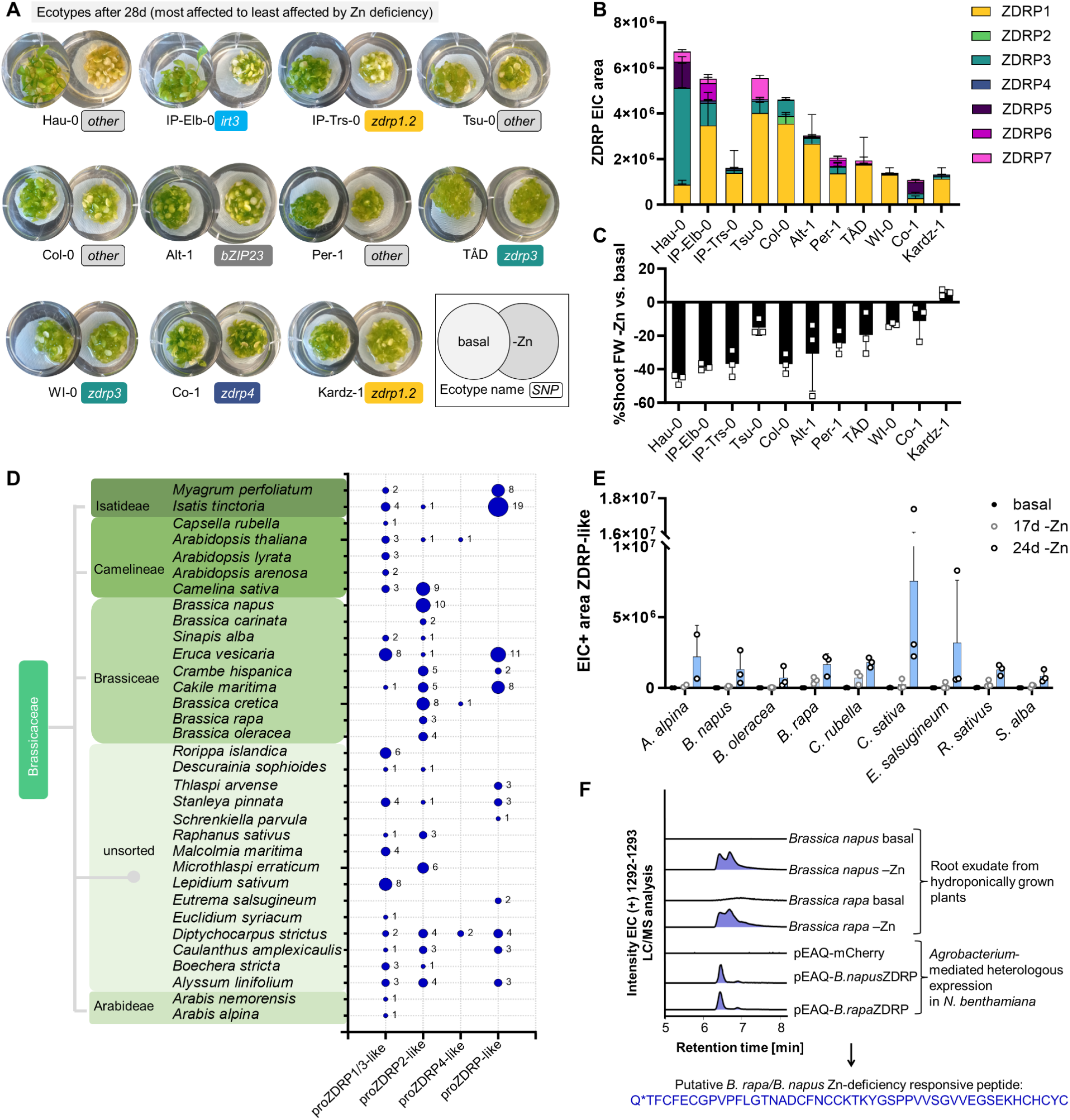
Distribution of ZDRPs among natural variants of *A. thaliana* and other Brassicaceae species. A) Phenotypes of natural variants under basal (+Zn) and -Zn conditions after 28 days. Selected SNPs indicated. Plants are ordered Zn-inefficient to Zn-efficient based on phenotypic comparison -Zn vs basal. B) LC/MS analysis of ecotypes root exudates grown under -Zn after 21 days. Ecotypes ordered by severity of Zn deficiency symptoms. Experiment performed in triplicate. Individual data points represent plants and root exudate in one well. Error bars represent standard deviations. C) Comparison of shoot fresh weight (FW) of plants grown under -Zn vs. basal conditions (-Zn growth compared to basal growth in %). D) Distribution of propeptide ZDRP amino acid sequences (proZDRP) predicted from Brassicaceae genomes. Circle size is proportional to occurrence of putative ZDRP sequences. E) LC/MS analysis of hydroponic media extract from Brassicaceae species grown under -Zn or basal conditions. Shown are the EIC values of the most abundant putative ZDRP per species. F) LC/MS analysis of *B. rapa*- and *B. napus*-derived ZDRPs in *N. benthamiana* after *Agrobacterium*-mediated transformation in comparison to root exudates of *B. rapa* and *B. napus* under basal and -Zn conditions. Q*, predicted pyroglutamic acid. Experiment performed in triplicate. Error bars represent standard deviations.

Following phenotypic assessment, we determined ZDRP levels in ecotypes. All natural variants produced a peptide response when grown under -Zn conditions. As predicted by the low SNP rate of *ZDRP1*.*1*, we noted that ZDRP1 was universally produced and could be detected as the predominant peptide in nine out of eleven ecotypes (Figure 3B). The higher number of SNPs in *ZDRP2-4* explains why ZDRP2-4 are not universally produced, with ZDRP2 made solely by ecotype Col-0. Intriguingly, root exudates of seven ecotypes revealed new mass features absent from Col-0 exudates: ZDRP5 (5361.4 Da), ZDRP6 (5350.3 Da), and ZDRP7 (5644.3 Da). The mass features of ZDRP5-7 were absent from +Zn root exudates coupled with their predicted size suggest they are peptides that are part of the ZDRP family in non-Col-0 ecotypes. Since their biosynthetic origin could involve a mutated *ZDRP* gene in non-Col-0 ecotype genomes, we investigated SNPs in selected ecotype genomes and found ZDRP7 to be a mutated version of ZDRP2 with an exchange of an arginine to a lysine (observed: 5644.3 Da, calculated: 5644.4 Da, Figure S7B-D).

When we compared yellowing of leaves and peptide production under -Zn conditions, we noted an apparent trend across the ecotypes (Figure S8). Hau-0, the ecotype most affected by Zn deficiency, produced the highest amount of ZDRPs while less affected strains like Kardz-1 produced the lowest ZDRP levels (Figure 3B). An inverse trend emerged when we compared plant growth under -Zn to basal conditions (basal/-Zn plant growth in %, Figure 3C). Aerial growth was most affected by ZDRP-overproducer Hau-0 under Zn deficiency (up to -45%) while less affected strains only lost about 20% of shoot weight under -Zn compared to basal conditions. In summary, by phenotypic assessment, metabolic profiling, and shoot weight variations we found a correlation between ZDRP production and plant health under Zn starvation. Furthermore, we discovered peptides that appear to expand the ZDRP family by three additional members absent from the model plant Col-0. Supported by the high conservation and production rate of geographically distant ecotypes, we found that ZDRP1 is especially crucial for low Zn response in *A. thaliana*.

Given all *A. thaliana* ecotypes produced a robust peptide response produced under -Zn conditions, we questioned whether ZDRPs are conserved among other genera within the Brassicaceae as well as other plant families. We first searched for ZDRP-homologous small protein sequences harboring one HCHC motif near the C terminus using the NCBI and the Phytozome database (42) in each candidate genome. Overall more than 25 species’ genomes harbor *ZDRP*-like genes, all of which were confined to the Brassicaceae. Hits include dietary plants like wild cabbage (*B. oleracea*), radish (*Raphanus sativus*), arugula (*Eruca vesicaria*), rapeseed (*Brassica napus*), and mustard (*Sinapis alba*) (Figure 3D and S9). Since Brassicaceae genomes have been sequenced extensively, we do not want to exclude the possibility of similar plant peptides encoded in more complex plant genomes. Initial screening metabolites in tomato (*Solanum lycopersicum*) and *N. benthamiana* did not reveal a similar peptide response when plants were grown under Zn deficiency (Figure S10).

To confirm production of Zn-responsive peptides in the Brassicaceae, we tested nine species in the hydroponics setup. Root exudates were collected for up to 45 days until phenotypic changes in the form of leaf yellowing in plants relative to those grown in Zn replete media was observed (Figure S9). Exudates were measured via LC/ESI-MS and the -Zn metabolic profile investigated for the occurrence of peptide masses absent from root exudates from +Zn media. These experiments revealed that putative ZDRP peptides are produced by all tested Brassicaceae under -Zn (Figure 3E). Brassicaceae peptides range between 5,000-7,300 Da although observed mass signatures do not match ZDRP1-4 (Figure S11). In fact, all species aside from *Brassica rapa, B. napus*, and *B. oleracea* produced their own unique set of Zn-responsive peptides. The absence of these Brassicaceae peptides from +Zn root exudates and their similar mass range make them possible ZDRP candidates to investigate in future studies.

As a proof-of-concept that candidate ZDRP-like sequences identified in the bioinformatic search correspond to detected Brassicaceae peptides, we selected a *B. rapa* gene also present in *B. napus* genome and employed *Agrobacterium*-mediated heterologous expression in *N. benthamiana*. The mass signature in pEAQ-*B*.*rapa* or pEAQ-*B*.*napus* infiltrated tobacco leaves matched the compound excreted from *B. rapa* and *B. napus* roots under Zn deficiency (Figure 3F). These data suggest that the HCHC motif can be used as search motif to identify related ZDRPs in land plant genomes. Notably, the peptides produced across these dietary Brassicaceae are heterogenous in sequence, however they are all actively produced when plants are cultivated under low Zn conditions.

### Peptide-deficient mutants, overexpression strains, and chemical complementation unveil a role for ZDRPs in root development under Zn deprivation

Defensins are well known for their toxicity, typically conferred by their cationic character that facilitates binding to membrane lipids (43). Several plant DEFL peptides have been shown to have strong antimicrobial activity and metal-binding properties (44). To explore if ZDRPs also target microbes, we tested ZDRPs against a panel of two fungal and four bacterial strains and monitored their growth rate with increasing peptide concentrations (6.25-100 μg/mL peptide concentration, SI methods) (Figure S12). No antimicrobial activity was observed in this experimental set-up which is consistent with the fact that transcript levels of ZDRP genes are specifically upregulated under abiotic and not biotic stress conditions (27, 29–33). Together, these data suggest ZDRPs true ecological role might involve more direct mechanisms to counteract Zn deficiency rather than eliminating potential competition for Zn, or accessing biotic micronutrient stores in the soil.

A role as zincophore (Zn-binding compounds) seemed a possible function for ZDRPs because small molecules that chelate Zn have previously been observed in bacteria and fungi (45, 46). Since ESI-MS leads to disposition of Zn in the stainless steel capillary rendering it unsuitable to investigate metal-binding of ZDRPs (47), we instead employed the softer ionization method MALDI-TOF MS (48) to determine if ZDRPs contain bound Zn. Typical amino acids for Zn coordination are histidine and cysteine. Since we detected ZDRPs in their oxidative form (disulfide bonds between cysteines), we first reduced ZDRPs. Applying tris(2-carboxyethyl)phosphine, TCEP, as a reducing agent to the whole root exudate led to a +8 Da shift corresponding to the reduced forms of ZDRP1-3 (Figure S13). When we added Zn^2+^ to the reduced peptide mix no mass shift was observed indicating a lack of binding (Figure S13). We also investigated putative Zn-binding capabilities of ZDRPs by Zincbindpredict (49), a tool to predict Zn binding sites, however, no typically Zn-binding sites were detected.

To investigate alternative ecological functions of ZDRPs such as in signaling and/or development, we looked for physiological or morphological changes in plants where ZDRP production is altered by T-DNA insertional mutation (50) or overexpression lines. Mutants of different parts of the Zn deficiency network that display impaired Zn responses relative to WT were chosen to investigate if ZDRPs could complement these genetic deficiencies and, consequently, reveal a functional role for ZDRPs. We included transcription factor mutants *bzip19* and *bzip23* as they have been shown to regulate plant responses to Zn deficiency (14) (Figure 4A). Zn transporter genes *ZIP3* and *ZIP4* were chosen based on their expression in -Zn transcriptomics datasets (27, 29–33), their dominant expression in root tissue (51), and their clear Zn response deficiency phenotypes. T-DNA mutants of *ZDRP* genes *zdrp1*.*1, zdrp1*.*2, zdrp2* were chosen for analysis since the ZDRP1 and ZDRP2 peptides are the dominant peptides secreted under Zn deficiency.

**Figure 4.**
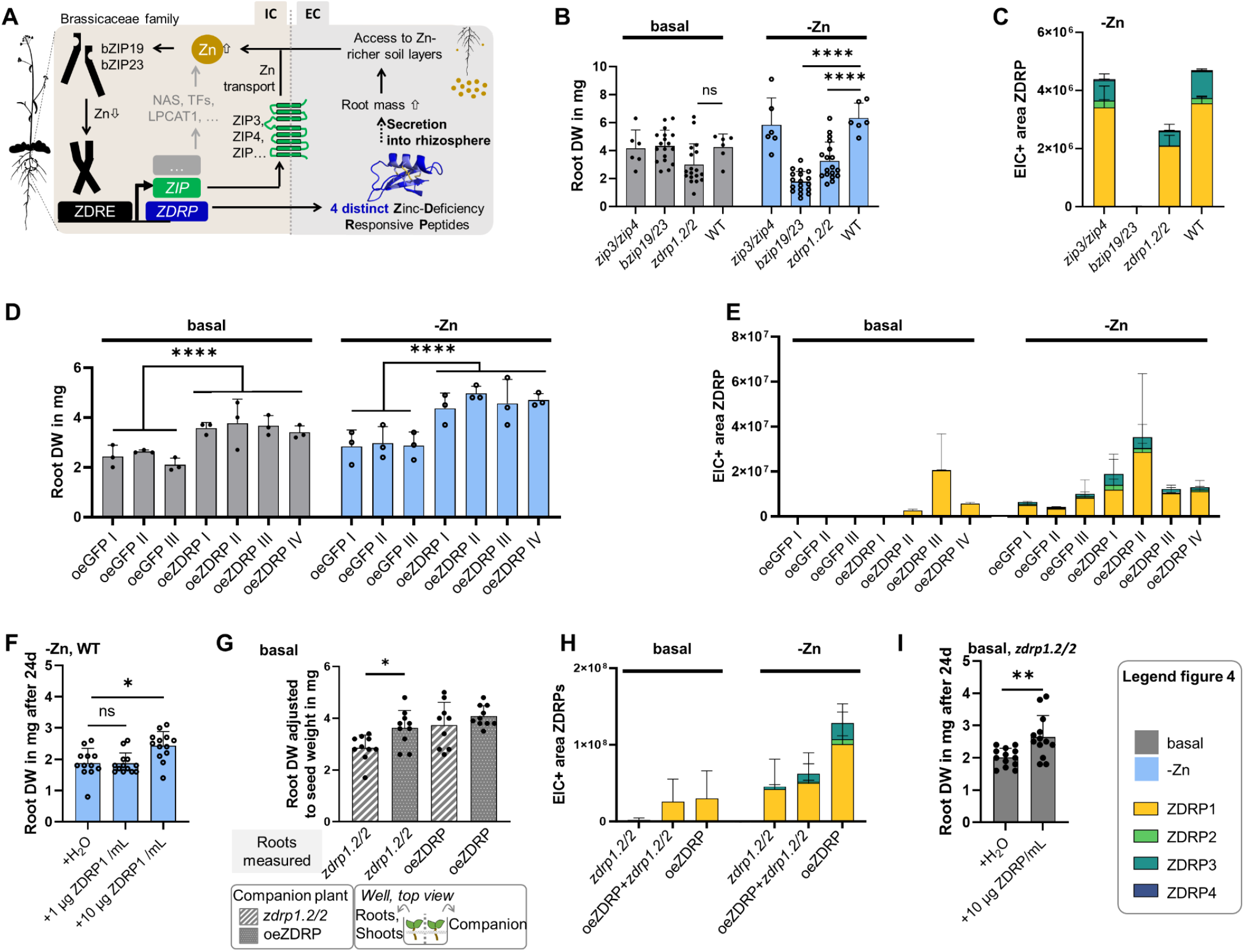
T-DNA mutants and overexpression lines unveil the ecological role of ZDRPs in root development. A) Overview of Zn-deficiency related network (bZIP, transcription factor; ZIP, zinc transporter; ZDRP, Zn-deficiency responsive peptides, TF, transcription factors, NAS, nicotianamine synthase). Plant adapted from bioicons. B) Root dry weights (DWs) of double mutants and WT grown hydroponically with or without Zn in the media after 28 days. C) LC/MS analysis of root exudates from double mutants and WT grown under Zn deprivation. D) Root DW of GFP-(oeGFP) and ZDRP-overexpression lines (oeZDRP) after 28 days. E) LC/MS analysis of root exudates from overexpression strains grown under basal conditions or Zn deprivation. Four (I-IV) independent lines. F) Root DW of WT plants after addition of vehicle control or ZDRP1 in -Zn media after 24 days. G) Root DW of WT and oeZDRP lines grown alone or next to a companion plant after 28 days. Values were adjusted to equal seed weight. H) LC/MS analysis of root exudates from plants shown in (G). I) Root DW of *zdrp1*.*2/2* plants after addition of vehicle control or ZDRPs in replete media after 24 days. Individual data points represent plants and root exudates in one well. One well has 4 mg of seed weight at the start of the experiments (ca. 250 seeds). Error bars represent standard deviations. Replicates per experiment: n=3-9 (4C), n=3 (4E), n=5 (4H). Statistical analysis, paired t-test. *P*-values: *, ⪬0.05; ***, ⪬0.0005; ****, ⪬0.00005.

Previous reports have revealed at least partial functional redundancy for transcription factors bZIP19 and bZIP23 in the evaluation of the individual gene mutants (14). We suspected that ZDRPs might also have some overlapping function. To address this possibility in a search for mutant phentoypes, we targeted the double T-DNA mutant lines *zip3/zip4, bzip19/23*, and *zdrp1*.*2/2*. All T-DNA double mutants were constructed by crossing of the respective single mutant lines and confirmed by PCR. These plants were indistinguishable in chlorophyll phenotype as well as shoot and root DW under basal conditions (Figure S14A-B). However, under Zn deficiency in hydroponic growth conditions, both the peptide mutant *zdrp1*.*2/2* and transcription factor *bZIP19/23* had a statistically significant loss in root dry weight (50 % and 67%, respectively) compared to the WT (Figure 4B). In contrast, shoot DW under Zn deficiency was comparable to WT for all mutants except *bZIP19/23* which showed a significant loss in shoot DW (Figure S14B). Mutant *zip3/zip4* performed similarily to WT plants indicating that any of the other ten ZIP transporters could be acting in a compensatory fashion in this genetic background.

We paired these growth phenotype analyses with measurement of ZDRP exudation. We observed a complete loss of all ZDRPs in the *bzip19/23* mutant; this finding corroborates previous transcriptomics data that indicate these transcription factors directly regulate ZDRP production (33) (Figure 4C). Single mutants *bzip19* and *bzip23* are still able to produce reduced, albeit measurable levels of ZDRPs (Figure S14C-D).

In contrast to the transcription factor mutants, we observed that the collection of *zdrp* single, double, and triple mutants produce varying levels of specific individuals ZDRPs. Metabolic profiling of root exudates of *zdrp1*.*2/2* showed no detectable ZDRP2 levels and reduced but detectable ZDRP1 for a net loss of 40% ZDRP levels relative to WT plants under -Zn conditions (Figure 4C). This is likely due to intact *ZDRP1*.*1* in the *zdrp1*.*2/2* mutant background. We wondered if the combination of two major secreted peptides – ZDRP1 and ZDRP2 – contribute additively to the root growth phenotype observed in WT plants under Zn deficiency. To test this, we next isolated a triple mutant *zdrp1*.*1/1*.*2/2* that we anticipated would be unable to produce both ZDRP1 and ZDRP2 (Figure S15A). Surprisingly, we found that *zdrp1*.*1/1*.*2/2* lost the root growth phenotype observed in *zdrp1*.*2/2* (Figure S15B). When monitoring ZDRP levels in *zdrp1*.*1/1*.*2/2*, we noted an significant increase in ZDRP3 levels in the root exudates comparable to whole ZDRP levels in the WT (Figure S15A). We speculate that the absence of ZDRP1 led to upregulation of *zdrp3*, and therefore accumulation of ZDRP3 peptide that is normally produced only at low levels in Zn-deficient WT plants or the *zdrp1*.*2/2* double mutant. Taken together, these data provide evidence that ZDRP gene products regulate root growth under Zn deficiency, and that different ZDRP peptides can each contribute to this phenotype. It is notable that ZDRP1 is highly conserved among the ecotypes, and it is not yet clear whether different ZDRPs lead to distinct root morphology changes that all contribute to a net increase in root mass.

We tested four independent transgenic lines (named oeZDRP I-IV) and generated GFP-overexpressing lines (named oeGFP) as a negative control (SI methods). We found that the ZDRP1-overexpression lines all showed higher root (but not shoot) mass relative to the oeGFP control. Furthermore, paired metabolite measurements revealed that the higher root masses corresponded to higher ZDRP production levels in these ZDRP1-overexpressing strains under both basal and -Zn conditions (Figure 4D-E, Figure S16). These results further support a model where secreted ZDRPs have a role in promoting plant root development specifically during Zn-deficiency stress.

In order to test the hypothesis that secreted ZDRPs and not other *zdrp* gene products have a direct role in root growth, we designed a chemical complementation experiment where purified ZDRPs are supplied exogenously to developing seedlings. To obtain a sufficient amount of peptide, we overexpressed ZDRP1 in *N. benthamiana* plants, and purified ZDRP1 from the apoplast extract using open-column chromatography followed by preparative HPLC purification. This method yielded 0.5 mg ZDRP1 from 15 *N. benthamiana* plants. In order to target biologically-relevant levels of peptide in complementation experiments, we determined that WT *Arabidopsis* seedlings produce ca. 3 μg ZDRPs/mL and oeZDRP IV up to 6 μg ZDRPs/mL in the absence of Zn after 28 days (based on standard curve in Figure S17A). To include this range of natural responses, we exogenously applied ZDRP1 to WT plants grown for 14 days under Zn-deficient conditions in two concentrations 1 μg/mL and 10 μg/mL (or 1.88 μM ZDRP1). We observed a significant increase in root mass, but not shoot mass, in plants grown in cultures supplemented with 10 μg ZDRP1 per mL medium (Figure 4F and Figure S17B). Increased ZDRP1 levels could be observed in root exudates of plants that were supplemented with ZDRP1 10 days post exposure revealing that added peptides stay stable in the medium over time (Figure S17C).

Results with exogenous addition of ZDRPs suggest roots can sense externally delivered peptide, however it remained unclear if the peptide signal can act in *trans* from a separate plant, how far ZDRPs migrate, and if they stay active in the hydroponic media after secretion. To address these questions, we grew *zdrp1*.*2/2* mutant and ZDRP1-overexpressing line alone or side-by-side in a hydroponics set-up. Shoot growth was not affected in *zdrp1*.*2/2* when grown side-by-side the ZDRP1-overexpression line in either basal or Zn-deficient conditions (Figure S18). However, we observed a significant increase in root weight in the *zdrp1*.*2/2* mutant when grown in the presence of the ZDRP1 overexpression line (relative to *zdrp1*.*2/2* grown alone) under basal conditions (Figure 4G). The same effect was not observed if the media was Zn deficient, likely due to ZDRP saturation from *zdrp1*.*2/2* cultures grown under -Zn conditions after 28 days. Replete media does not induce ZDRP production in *zdrp1*.*2/2* (Figure 4H). Thus, addition of ZDRP1 supplied by ZDRP1-overexpressing lines to *zdrp1*.*2/2* cultures grown in replete medium induces extensive root growth promoting effects. ZDRP levels corresponded linearly with the absence or presence of the ZDRP1-overexpressing strains and of Zn (Figure 4H).

To corroborate that ZDRP addition to Zn-containing basal medium can influence plant root growth, we investigated root DW of double mutant *zdrp1*.*2/2* grown in replete medium either in the presence of 10 μg ZDRPs per mL medium or vehicle control (Figure 4I). We observed a significant increase in root DW of plants supplemented with ZDRP levels when compared to no supplementation after 24 days. This finding is also in coherence with the observation that overexpression lines of ZDRP1 show enhanced root growth even in replete media.

In summary, these results strongly suggest that ZDRPs exuded from roots stay active in the media over time, and can function as an autocrine signal between plants to increase root growth. It is possible that an expanded root architecture could give access to soil layers that are potentially Zn-richer which is advantageous for plant survival.

## Discussion

In this study we uncovered a new class of DEFL peptides, named Zinc-Deficiency Responsive Peptides (ZDRPs), that are actively secreted by plant roots and result in root remodeling under Zn deficiency.

DEFL peptides are related to defensins, a class of peptides from animals, plants, and insects (26). There are over 300 predicted *DEFL* peptide genes with mostly elusive products and ecological roles that appear to be outside of defense (26). DEFL peptides have been known to function in immunity (52) and, in recent years, have been found to be involved in determining self-incompatibility (53), in pollen development (54), in plant-microbe symbiosis (55) and in tolerance to metal toxicity (56). ZDRP screening against a variety of fungal and bacterial pathogens did not reveal an antimicrobial effect. Similarly, a previous study found only minor antinematodal effects of ZDRP1 (15). *ZDRP1*.*1* expression has been monitored using a GUS fusion in *Arabidopsis*, and has been shown to be exclusive to roots and enhanced in roots infected with nematodes (15). As nematodes and plants fight for valuable nutrients in the soil (57), it is feasible that nematode infection could have triggered Zn-deficiency response in form of ZDRPs. Since *ZDRP1-4* expression is controlled by bZIP19 and bZIP23, their expression has a clear connection with abiotic stress responses (Zn deficiency) rather than a singular role in biotic stress. The addition of a Zn deficiency response to the diverse list of ecological DEFL roles expands our understanding of this peptide family.

A recent report focused on the role of enhanced gene expression of *ZDRP2* (*DEFL203*) and *ZDRP4* (*DEFL202*) under Zn deficiency in *A. thaliana* (16). In this study it was found that *zdrp2/4* double mutant led to enhanced root meristem length and cell number compared to WT plants. However, root changes in response to Zn status were not observed when grown on agar in *ZDRP4* overexpression lines and mutants relative to WT (16). In our study using an alternative hydroponic model for growth, we observe a root growth-promoting effect of the peptide ZDRP1 that is supported by chemical complementation, *in trans* complementation, and four independent overexpression lines. We find that ZDRP1 is the most conserved and mainly produced peptide by Brassicaceae under Zn deficiency, and therefore concentrated our investigations on this member of the CRP0770 family. Since the primary peptide sequences of ZDRP2 and ZDRP4 vary from ZDRP1 and ZDRP3, it is possible that each peptide might have a different effect on the root architecture and development. Future studies will shed light on the various effects of ZDRPs.

Several different peptides triggered by the plant nutrient status are known. They involve peptides from the CLAVATA3/EMBRYO SURROUNDING REGION-RELATED (CLE), C-TERMINALLY ENCODED PEPTIDE (CEP), and ROOT GROWTH FACTOR (RGF)/GOLVEN (GLV)/CLE-Like (CLEL) family (58). CEP1 is produced under nitrogen starvation, modulates root development and enhances nutrient uptake of inorganic nitrogen from the soil (59–61). CLE1, CLE3, CLE4, and CLE7 serve as a root-to-shoot signal leading to an expanding root system under low nitrogen levels (62). CLE14, a positive regulator of root hair formation (63, 64), and RGF1-3, which control vertical root growth, are produced under phosphate deprivation (65–67). All of these previously described peptides serve a role under macronutrient but not micronutrient deficiency. Their core peptide sequence of CLEs and RGFs is 12-15 amino acid long. In contrast, ZDRPs represent typical DEFL peptides with a core sequence of 49-51 amino acids. Although exogenously application of non-physiological amounts of peptide can lead to a non-specific response in plants, these effects are defined by reduction in root development as seen for CLE26 (68). Contrary to this, we observe a positive influence on root mass under ZDRP treatment and in overexpression lines. ZDRP size, their specific production under micronutrient stress, and their putative positive involvement in root growth promotion distinguish them from these previously known examples.

There are two examples of peptides that share functional similarities to ZDRPs which might suggest ZDRP possible mode of action: IRONMAN (IMA) peptides that respond to the Fe status of the plant (69, 70) and RALF peptides. IMA peptides require a specific motif to exert their role in iron homeostasis, however their sizes are highly variable, ranging from 23-86 amino acids (70). Overexpression of IMA peptides leads to upregulation of Fe uptake genes in the roots of angiosperms. Even though ZDRPs and IMA peptides are produced under micronutrient stress, their structure and ecological roles vary significantly. Based on absence of posttranslational modifications, peptide size, and a function in root development, ZDRPs appear more closely related to the 5 kDa RALF peptides. In addition to their role in fertilization (71), RALF peptides influence root development negatively through auxin induction (72) but are essential for root hair formation (73). In contrast, we observe positive influence of ZDRPs on root development, suggesting a possible alternative mechanism of action.

Enhanced root development mediated by peptide hormone signalling under nutrient deficiency has been observed in plants (1). If plants are lacking the essential macronutrient nitrogen, the CEP/CEPD (C-TERMINALLY ENCODED PEPTIDE DOWNSTREAM) pair leads to enhanced root growth. For Zn deficiency, we found a parallel in ZDRPs as they are produced by plants to access Zn-rich soil levels.

Here we added another piece to our understanding of how plants combat Zn starvation. Transcription factors bZIP19 and bZIP23 form dimers under low Zn and bind to ZDRE motifs in promoter sequences of target genes including five *ZDRP* genes coding for the production of 5-6 kDa peptides in Brassicaceae (Figure 4A). After cleavage of the signal peptide sequence, ZDRPs are secreted into the rhizosphere where they act as autocrine signals to promote root growth. ZDRPs are part of the exploratory strategy of Brassicaceae to reach Zn-richer deposits in the soil. These findings will be useful in future engineering approaches to make plants more resilient to growth in low Zn conditions as more and more arable land becomes Zn deficient through natural causes and extensive application of fertilizers.

## Material and Methods

### Material availability

Double mutants and triple mutant will be purchasable from ABRC. Overexpression plasmids will be made available at ABRC.

### Methods

Methods are available in the Supporting Information online.

## Acknowledgements

We thank MOI lab at University of California Berkeley for providing ecotype seeds and helpful discussion, Paul Hoang for technical assistance, and Vivian Zhong (Brophy lab, Stanford University) for providing advice. *Brassciaceae* seeds were a gift from the Dinneny lab (Stanford University). MALDI-TOF analysis was performed by the PAN facility (Stanford University, Jessica Tran). We thank Undramaa Bat-Erdene for helpful discussion. This work was funded by the Deutsche Forschungsgemeinschaft (DFG, German Research Foundation) – project no. 516932270 (to S.P.N.).

