## Supplemental Information for "Zinc-starved Brassicaceae Plants Secrete Peptides that Induce Root Expansion"

**This PDF file includes:**

Experimental Procedures

Tables S1 to S4

Figures S1 to S18

Supplemental References

### Experimental Procedures

##### Plant material.

Unless stated otherwise, all *Arabidopsis thaliana* ecotypes and *Nicotiana benthamiana* were grown under standard greenhouse conditions (16:8 photoperiod at 130 µmol/m$^2$/s, 21 °C constant temperature and 50% relative humidity). PRO-MIX BX (Premier Tech, USA) fertilized with Osmocote 14-14-14 (ICL, USA) was used as a growing substrate.
*Eutrema salsugineum* PRS 69 var. Shandong and *Capsella rubella* PRS 224 seeds were a gift from José Dinneny (Stanford University). *Arabis alpina* (White Rockcress) seeds were purchased from Seedville (USA). *Brassica oleraceae* seeds were purchased from Johnny's Selected seeds (USA). The seeds of the remaining Brassicaceae family were purchased from Strictly Medicinal seeds (USA). *A. thaliana* ecotypes were a gift from Moises Exposito-Alonso (UC Berkeley, USA).
The following T-DNA lines were obtained from ABRC: SALK_008709 (AT2G32270, *ZIP3*), SALK_145371C (At1g10970, *ZIP4*), SAIL_520_G01 (At5g33355, *ZDRP1.2*), SAIL_1251_D11 (At2g36255, *ZDRP2*), SALK_018248C (At2g16770, *bZIP23*), SALK_144252C (At4g35040, *bZIP19*), SAIL_628_B05 (At3g59930, *ZDRP1.1*). The mutants were confirmed by PCR with GoTaq® DNA Polymerase (Promega) from genomic DNA. Oligos were designed by T-DNA Primer Design tool from SIGnAL (http://signal.salk.edu/tdnaprimers.2.html) (Table S1). All genotypes were confirmed as homozygous by PCR in successive generations (4$^{th}$ generation).

##### Seed sterilization and stratification.

Seeds were washed first with 70% ethanol for 40 seconds. *A. thaliana* seeds were sterilized with 1.5% sodium hypochlorite solution and 0.2% Tween 20. Other Brassicaceae seeds were sterilized by adding 3% sodium hypochlorite with 0.2% Tween 20. Seeds were thoroughly washed with sterile water and stratified for 2-3 days at 4 °C in the dark before the start of the assays.

##### Zinc-deficiency assay (hydroponics).

Small scale *A. thaliana* var. Col-0 experiments were performed in sterile 6-well plates (Corning). A rigid PTFE plastic mesh with a 0.025" x 0.005" opening and 12" wide (McMaster-Carr) was cut to fit into the wells and sterilized in a separate container. Three milliliters of sterile Murashige and Skoog medium (MS, Table S2, pH 5.7) with (basal) or without zinc (-Zn) were added to the corresponding well. Four mg of seeds were added to each well. The well plates were sealed with microporous tape (3M). Medium was exchanged after 14 days and 21 days. The assay stopped on day 28. Plants were grown at 22 °C with constant light.
For the Brassicaceae screen, plants were grown on floating rafts in magenta boxes containing 20 ml of basal or -Zn medium per box. All Brassicaceae seeds except for *E. salsugineum, A. alpina,* and *A. thaliana*, were grown on PTFE plastic mesh with a 0.04" x 0.025" opening and 18" wide (McMaster-Carr). *E. salsugineum, A. alpina,* and *A. thaliana* were grown on the same mesh used for *Arabidopsis* mentioned above. Due to the size differences of all Brassicaceae seeds, different number of seeds were placed in each magenta box: 6 seeds for *Camlina sativa*; 10 seeds for *Sinapis alba, Brassica napus, Brassica rapa, Raphanus sativus, Brassica oleraceae*; or, 20 seeds for *E. salugineum* or *A. alpina.* The media was exchanged after 17 days, 24 days, 31 days, and the assay ended at day 45.
For the *S. lycopersicum* vf36 and *N. benthamiana* screen, plants were grown on floating rafts in magenta boxes containing 20 mL of basal or -Zn medium per box. All seeds were grown on PTFE plastic mesh with a 0.04" x 0.025" opening and 18" wide (McMaster-Carr). We sowed 10 sterile and stratified seeds per box. The media was exchanged after 7 days and 14 days. The assay ended at day 21.
The relationship between chlorophyll content, zinc(II) concentration and ZDRP production was determined using a small-scale setup: Hydroponics studies were performed in 6 well plates with 4 mg seeds per treatment with decreasing zinc concentrations (100%, 50%, 20%, 10%, 5%, 2.5%, 1%, 0.2%, 0%) where 100% zinc corresponded to 2.9 µM $ZnSO_4$. Medium was exchanged after 14 days and 21 days.

To evaluate if re-supply of zinc(II) abolishes ZDRP production, the following set-up was used: Four mg of sterilized and stratified *A. thaliana* var. Col-0 seeds were germinated hydroponically in the absence of zinc in one well of a 6-well plate. After 14 days, spent media was replaced. Seven days after replacing the media (day 21), root exudates were collected, and media was substituted with new media with (basal) or without zinc (-Zn). Root exudates and plant tissues (roots and shoots) were collected on days 26, 28, and 31. Aerial tissue was also used for chlorophyll quantification.
Root exudates were collected as 1.5 mL fractions in 2 mL tubes, frozen at -80 °C, and lyophilized to dryness for further analysis.
To determine if closeness to ZDRP1-overexpressing lines can induce increased root weight in the *zdrp1.2/2* mutant, we seeded either 4 mg of plant lines alone on PTFE mesh per well or 2 mg of *zdrp1.2/2* mutant on a PTFE mesh floating next to 2 mg of oeZDRP1 seeds floating on a separated PTFE mesh. Hydroponic experiments were performed as indicated under small-scale *A. thaliana* experiments.

**Metabolite extraction.**

Root exudates were collected from hydroponic growth experiments, frozen, and lyophilized. 60 µL 80% methanol in water per mL root exudate was added to the crude extracts and vortexed. The mixtures were sonicated for 15 min, centrifuged at 10,000xg for 5 min, and the filtered supernatant was used for LC/MS analysis.
Plant tissue was frozen in liquid nitrogen and lyophilized to dryness. Tissue was homogenized on a ball mill (Retsch MM 400) using 5 mm diameter stainless steel beads, shaking at 25 Hz for 2 min at room temperature. Tissue powder was resuspended in 30 µL 80% methanol per mg dry weight, centrifuged at 10,000 rpm for 5 min, and measured by LC/MS.

**Root and shoot measurement.**

Roots below and shoots above the PTFE mesh were cut by a razor blade, washed in Millipore water, and transferred to pre-weighted 2 mL or 5 mL tubes for roots and shoots, respectively. Tissues were dried at 60 °C for 2 days or lyophilized for 24 hours. Dry weight (DW) was determined with a precision balance.

**Chlorophyll measurement.**

Lyophilized shoot tissue was homogenized on a ball mill (Retsch MM 400) using 5 mm diameter stainless steel beads, at a frequency of 25 Hz for 2 min. Metabolites were extracted with 100% methanol as 1 mL per 10 mg DW. Samples were centrifuged at 10,000xg at room temperature for 5 min. The absorbance of 1:10 dilutions in methanol was determined spectrophotometrically at $\lambda = 666$ nm and $\lambda = 653$ nm. Chlorophyll content was calculated using the equations developed by Wellburn (1). All measurements were performed at 20 °C using disposable plastic cuvettes on a Genesys 20 spectrophotometer (ThermoFisher).

**ZDRP secretion in soil.**

*A. thaliana* var. Col-0 seeds were grown on half-strength MS agar (0.8% agar, pH 5.7) for three weeks. At that point, the effects that pH, soil composition and permeability have on ZDRP secretion were determined. To test the effect of pH, Pro-Mix BX pH was adjusted to pH 7, pH 8, or pH 10.6 using 6 M NaOH. To test the effect of soil composition and permeability, two alternative approaches were carried out. In the first approach, Pro-Mix BX (composition: 75-85% sphagnum peat moss, perlite, limestone, wetting agent, mycorrhizae) was mixed with sand (Mg Al Fe silicate, Specialty Vermiculite Group) in the following ratios: 1:1, 1:2, 1:3, 1:4, 1:5, 1:6, 1:7. On the second approach, Perlite (MiracleGo) and sand was mixed 1:1 or 1:2 and watered with 50 mL per pot of MS medium (without sucrose and without Zn) was added to the mixture. In all cases, 5 plants were grown in each pot. Soil conditions were tested in duplicates.

After 6 weeks 5 mL of soil was taken from the rhizosphere of the surviving plants and 5 mL of water were added to the soil sample, centrifuged at 4,000 xg for 10 min and 3.5 mL of the supernatant were lyophilized. The obtained solids were dissolved in 200 µL of 80% methanol, ultrasonicated, centrifuged at 4,000 xg for 5 min and the supernatant filtered (0.45 µm low-binding PTFE, Millipore). The samples were analyzed by LC/MS (Figure S1G).

**Cloning.**

All oligos for cloning included overhangs to allow for Gibson Assembly to the appropriate vector (pEAQ for transient, pYB1301 (1301Y) for stable expression; see Table S3).

For transient expression we employed pEAQ-HT as a backbone (2). The pEAQ-HT plasmid was digested with XhoI and AgeI following the manufacturer's instructions (NEB). Genes of interest were PCR-amplified from *A. thaliana* genomic DNA or *Brassica rapa* or *Brassica napus* gDNA. Digested pEAQ-HT and amplified PCR products were assembled using NEBuilder HiFi DNA Assembly Mix (NEB). Plasmids were transformed into *E. coli* NEB DH10-beta (Selection 50 µg/mL of kanamycin). Plasmid DNA was transformed into electrocompetent *A. tumefaciens* GV3101. Selection was carried out at 30 °C for 2d using LB agar plates containing 50 µg/mL kanamycin and 30 µg/mL gentamicin.

For stable expression, we employed pYB1301 (1301Y) as a backbone (3). The plasmid was digested with PmeI and PmlI according to the manufacturer's recommendations (NEB). Gene *ZDRP1.2* (At5g33355) was PCR-amplified from *A. thaliana* gDNA (Table S3) using Q5 High Fidelity DNA Polymerase (NEB) introducing overhangs. Digested 1301Y and amplified PCR products were assembled using homologous yeast recombination. The plasmid was introduced into chemically competent *E. coli* 10-β (selection 50 µg/mL kanamycin), isolated, sequenced, and introduced into electrocompetent *A. tumefaciens* GV3101 (selection 50 µg/mL kanamycin, 30 µg/mL gentamicin).

**Heterologous expression of ZDRPs in *N. benthamiana*.**

*Agrobacterium tumefaciens* cells were grown in LB medium containing 50 µg/mL of kanamycin and 30 µg/mL gentamicin at 30 °C for 2 days. The cells were centrifuged at 3500xg for 5 min. The pellet was resuspended in *Agrobacterium* induction medium (10 mM MES pH 5.6, 150 µM acetosyringone, 10 mM $MgCl_2$) and incubated at room temperature for 2 h. *A. tumefaciens* suspensions were prepared at $OD_{600nm}$ 0.2 and infiltrated in 5-week-old *N. benthamiana* plant leaves using a needleless 1 mL syringe. Leaves were collected after 5 days of growth, frozen in liquid nitrogen, grinded with stainless steel beads for 2 min at 25 Hz, and resuspended in 50 µL 80% methanol per mg plant tissue. The filtered samples (0.45 µm low-binding PTFE, Millipore) were measured by LC/MS.

**Apoplast extraction**.

*Agrobacterium*-infiltrated leaves were collected after 5 days, submerged in buffer solution (20 mM MES pH 6.0, 2 mM $CaCl_2$, 0.1 M NaCl) and vacuum infiltrated for 5 min. Leaves were rolled on Parafilm and introduced into the barrel of 5 mL syringes that hung from the flange in a 15 mL conical tube. The apoplast extract was obtained by centrifugation at 4,000xg for 10 min at room temperature. Apoplast extract was filtered (0.45 µm low-binding PTFE, Millipore) and diluted 1:1 in 100% methanol prior to LC/MS analysis.

**Generation of overexpression strains.**

We applied 1301Y-GFP as negative control. Plasmid 1301Y-ZDRP1 was employed to generate the ZDRP1-overexpressing strain.

*A. thaliana* Col-0 were transformed by the flower dip method (4). Flower dipping was repeated after 2 weeks. Harvested seeds were sterilized, stratified at 4 ℃ for 3 days, and seeded on ½ MS agar (0.8% agar, pH 5.7) supplemented with 50 µg/mL hygromycin. The plants were grown under standard greenhouse conditions. The presence of the introduced *ZDRP1.2* under the control of the 35S promoter in the hygromycin B-resistant seedlings was investigated by PCR using GoTaq Polymerase (Promega) and oligos SN13 and SN14 (Table S3).

To confirm that plants produce ZDRPs, a subset of plants were grown for 7 days in 6 well plates containing MS medium lacking zinc (1 plant per well floating on PTFE mesh in 3 mL medium). The root exudate in each well was collected, lyophilized, and resuspended in 90 µL 80% methanol, ultrasonicated for 15 min, centrifuged at 10,000 rpm for 5 min, and the filtered supernatant measured by LC/MS. In addition, leaf material from the same plants was collected, lyophilized, ground with stainless steel beads as mentioned above, and resuspended in 100 µL 80% methanol per mg plant tissue. The filtered samples were measured by LC/MS.

**Bioinformatics analysis.**

**Phylogenetic tree.** We performed a similarity search of the ZDRP1 amino acid sequence against the NCBI and Phytozome database (5) and selected the top100 hits. MEGA11 (6) was employed for aligning the sequences using the ClustalW method (7). We excluded sequences that were missing the characteristic HCHC motif as well as truncated protein sequences. A phylogenetic tree was constructed as maximum likelihood tree using MEGA11. Bootstrapping was set at 100. The phylogenetic trees were rooted to outgroup *Arabidopsis halleri* DEFL CCN97888.1. Examples of ZDRPs are listed in Table S4.
**SignalP (server27).** Predicting the cleavage sites between signal and core peptide was achieved by using SignalP v 6.0 (8).
**AlphaFold.** Protein structure prediction was performed by using AlphaFold (9). Uniprot ID for ZDRPs used: Q3E8R5 (ZDRP1), Q2V424 (ZDRP2), Q2V2Q8 (ZDRP3), Q2V3J6 (ZDRP4). For modeling we used PyMOL. The signal sequence was omitted and predicted disulfide bridges highlighted.
**HMMsearch.** HMMsearch (HmmerWeb version 2.41.2) (10) was performed to search against the default reference proteome database.
**Polymorph1001.** Polymorph1001 (11) was used to identify mutation rate of Zn-related genes in *A. thaliana* ecotypes.
**ScanProSite** (12) was used to identify amino acid pattern of ZDRP1 and ZDRP2 based on MS/MS results.
**Pattern Locator** (13) was used to identify ZDRE motifs in the promoter sequences of *ZDRP1-4*.

**Statistical analysis**.

Data was analyzed using unpaired, two-tailed Student's t-test. All experiments were repeated at least twice with ≥3 replicates.

**Antimicrobial activity assays.**

To assess the antifungal activity of ZDRPs on *Botrytis cinerea* SF1 and *Alternaria brassicicola* FSU218, 100 µL of a 2:100 dilution in PDB of a peptide solution in DMSO was added in each well of a 96 well plate containing $2x10^4$ spores/mL in 2xPDB pre-germinated for 7 hours in the dark. Plates were sealed with Microporous tape (3M) and $OD_{600}$ was recorded at regular intervals for 72 hours on a VersaMax microplate reader (Molecular Devices).
For antibacterial assays, various bacterial strains (*Escherichia coli* TOP10, *Pseudomonas syringae* pv. *maculicola* ES4326, *Agrobacterium tumefaciens* GV3101, *Bacillus subtilis* EG219) were pre-cultured in LB medium at 30 °C for 10 hours and diluted to $OD_{600}$ 0.1. Ten µL of bacterial suspension was added to a 96-well microtiter plate containing 87.5 µL of LB medium and 2.5 µL of ZDRP mix. Growth at 30 °C was monitored on a plate reader by measuring absorbance at 600 nm for 24 hours. The protein content of ZDRP mix was estimated by NanoDrop 1000 spectrophotometer at 280 nm normalized to BSA.

**Chemical complementation with ZDRP1.**

Peptide purification was carried out as described in "**Purification of ZDRP1**" below. ZDRP1 solution (1 mg/mL) was prepared in Millipore water and sterile filtered. *A. thaliana* var. Col-0 seeds were grown in a hydroponics set-up as previously described. Four mg seeds were added to each well of a 6-well plate. The plants were grown under -Zn conditions for 14 days. The medium was exchanged with water, 1 µg ZDRP1 per mL or 10 µg ZDRP1 per mL to the respective wells, and plants were grown for 10 more days. At the end of the experiment, root exudates, roots, and shoots were harvested. The tissues and exudates were lyophilized and analyzed by LC/MS. The chlorophyll content was determined as described above.

#### General analytical methods

##### LC/MS analysis.

**C18, reversed phase.** Metabolite analysis was performed on an Agilent 1260 HPLC coupled to a 6520 Q-TOF MS, using a 5 µm, 2 mm × 100 mm Gemini NX-C18 column (Phenomenex). Injection volume was 3 µL. The ionization energy employed was 150V. Skimmer 65V, OctIRFVpp 750 V, 325 °C gas temperature, 10 L/min drying gas, nebulizer 35 psi, mass range: 50-1,700 Da. Flow rate was 0.4 mL/min. Mobile phases were water supplemented with 0.1% formic acid (A) and acetonitrile supplemented with 0.1% formic acid (B). Gradient: 0–1 min 3% B; 1–10 min, 3–50% B; 10–13 min, 50–97% B; 13–15 min, 97% B; 15–15.5 min, 97–3% B; 15.5–21 min, 3% B.

**HILIC.** Metabolite analysis was performed on an Agilent 1260 HPLC coupled to a 6520 Q-TOF MS, using a 2.7 µm, 1.2 mm × 100 mm HILIC-Z Poroshell 120 column (Agilent). Injection volume was 3 µL. The ionization energy employed was 150V. Skimmer 65V, OctIRFVpp 750 V, 325 °C gas temperature, 10 L/min drying gas, nebulizer 35 psi, mass range: 50-1,700 Da. Flow rate was 0.25 mL/min. Mobile phases were 10 mM ammonium formate supplemented with 0.1% formic acid (A) and 10 mM ammonium formate in 90% acetonitrile (B). Gradient: 0–3 min 100% B; 3–13 min, 100–60% B; 13–14 min, 60–100% B; 14–20 min, 100% B.

**MS/MS fragmentation.** For MS fragmentation, 35 V was employed as collision energies with an *m/z* window of +/- 1.3.

##### Purification of ZDRP1.

Large scale expression and purification of heterologous expressed ZDRP1 was accomplished by infiltrating 15 plants following an up-scaled version of *N. benthamiana* transient Agroinfiltration. Briefly, 50 mL of *A. tumefaciens* GV3101 carrying pEAQ-ZDRP1 was grown as described above, induced with acetosyringone and resuspended to an $OD_{600}$ 0.5 in 3 L induction medium (20 mM MES pH 6.0, 2 mM $CaCl_2$, 0.1 M NaCl). Whole 5-week old tobacco plants were infiltrated by submerging the aerial tissue in a beaker containing 1 L of bacterial suspension and subjecting the plants to 600 mm Hg vacuum for 3 min for 3 times. Complete infiltration was evidenced by waterlogging of leaves. Plants recovered rapidly under standard greenhouse conditions and were grown for an additional 5 days. Apoplast washes from 72 infiltrated leaves were obtained, frozen and lyophilized.

Alternatively, we obtained ZDRPs from the native host by carrying out a large-scale *A. thaliana* cultivation according to the protocol developed by Monte-Bello *et al.* (14). We seeded *A. thaliana* var. Col-0 in 0.5x MS agar (1% agar, without sucrose, pH 5.7) and filled p200 sterile pipette boxes with 0.5x MS medium without Zn. After 6 weeks, the root exudates were harvested and lyophilized. The crude extract was resuspended in 4 mL distilled water. All ZDRP solutions were filter sterilized through a 0.22 µm filter before use.

A SPE column (Sep-Pak Vac, 12cc, 2 g, C18 cartridge, Waters) was preconditioned by washing with 8 mL methanol followed by 8 mL distilled water, before loading with the apoplast extract. After loading, the column was washed with an additional 9 mL of water before eluting the peptides with 4 mL of methanol. The final purification was carried out on a C18 column (CLIPEUS, 5 µm, 250 x 10 mm, Higgins Analytical INC.) in a preparative HPLC operating at a flow rate of 5 mL/min and an injection volume of 400 µL. Mobile phases: distilled water with 0.1% formic acid (A) and acetonitrile with 0.1% formic acid (B). Gradient: 0-0.5 min 5% B, 0.5-12 min 5-50% B. The absorbance was monitored at 280 nm and 254 nm.

##### Peptide reduction and investigating zinc binding via MALDI-TOF analysis.

Three mL root exudates were harvested from 4 mg of seeds of overexpression strain oeZDRP IV grown under -Zn conditions after 28 days and lyophilized to dryness. Crude extract was resuspended in 40 µL of Millipore water and split into three 1.5 mL microtubes. Total protein concentration was estimated as >10 mg/mL by absorbance at 280 nm. For peptide reduction, we added 6.5 µL of 10 mM TCEP solution (pH 7.0, final concentration 3.3 mM TCEP). To assess zinc binding capabilities, we added 6.0 µL of 10 mM TCEP solution (pH 7.0, final concentration 3 mM TCEP) and 1 µL 100 µM Zn(II) solution (pH 7.0, final concentration 5 µM Zn(II) solution). In all cases, MilliQ water was added to a final volume of 20 µL. Samples were incubated at 60 °C for 35 min. The reactions were stopped by the addition of 1:10 10% TFA.

Samples were cleaned using zip tip C18 10 µL tips according to the manufacturer's recommendations (Millipore). All cleaning steps were performed in the presence of 0.1% TFA. Final peptide elution was performed in 50% acetonitrile+0.1% TFA. Peptide solution was spotted together with either DHB (2,5-dihydroxybenzoic acid) or SA matrix (sinapinic acid) on a grid and analyzed on a Perseptive Biosystems (ABI), Voyager-DE RP, MALDI-TOF mass spectrometer equipped with Delayed Extraction technology (DE).

### Supplemental Tables

**Table S1.** Oligos used in this study to confirm T-DNA lines.

| **Primer name** | **Sequence 5'→3'** | **Notes** |
|---|---|---|
| SAIL_1251_D11_LP<br>SAIL_1251_D11_RP | TCAGCTACATTTACGGTGCTG<br>TTTTTGGCGGAAAAGCTAATC | T-DNA confirmation primers for *zdrp1.2* (At5g33355) |
| SAIL_628_B05_LP<br>SAIL_628_B05_RP | TATCTTCGGAAACGGATGTTG<br>CGCAAAGCATTAAAGAAATGG | T-DNA confirmation primers for *zdrp1.1* (At3g59930) |
| SAIL_520_G01_LP<br>SAIL_520_G01_RP | TTCCTCCAAAAGGTTGGAAAG<br>AAAAAGTTGCCAAGATTCGTG | T-DNA confirmation primers for *zdrp2* (At2g36255) |
| SALK_008709_LP<br>SALK_008709_RP | CCTTTAATGAAAGCTCTATTTCCG<br>GATGTTGTCTTTGTCGAAGCC | T-DNA confirmation primers (*zip3*) |
| SALK_145371_LP<br>SALK_145371_RP | AGATTGCTGTTGTTCCTGTGG<br>AATGTTGCCCTAATTTGGCTC | T-DNA confirmation primers (*zip4*) |
| SALK_018248_LP<br>SALK_018248_RP | TCTCTTATTAACGCCGGATCC<br>TGCAATTCGAAATTCCTTTTG | T-DNA confirmation primers (*bZIP23*) |
| SALK_144252_LP<br>SALK_144252_RP | ATTGACGTTGCTGAATGATCC<br>ACGATGCCATCTGTTTAGTGC | T-DNA confirmation primers (*bZIP19*) |
| LBb1.3<br>LB1 | ATTTTGCCGATTTCGGAAC<br>GCCTTTTCAGAAATGGATAAATAGCCTTGCTTCC | LBb1.3 for SALK lines,<br>LB1 for SAIL lines,<br>T-DNA confirmation primers |

**Table S2.** Modified MS medium components (15).

| Compound | Amount g/L | Stock solution |
|---|---|---|
| Sucrose | 5 | macronutrient |
| Potassium nitrate | 1.9 | macronutrient |
| Ammonium nitrate | 1.65 | macronutrient |
| MES | 0.5 | macronutrient |
| Calcium chloride dihydrate | 0.44 | macronutrient |
| Magnesium sulfate heptahydrate | 0.37 | macronutrient |
| Potassium phosphate, monobasic | 0.17 | macronutrient |
| Boric acid | 0.0062 | Micronutrient/vitamin |
| Cobalt chloride hexahydrate | 0.000025 | Micronutrient/vitamin |
| Cupric sulfate pentahydrate | 0.000025 | Micronutrient/vitamin |
| Manganese sulfate monohydrate | 0.0169 | Micronutrient/vitamin |
| Sodium molybdate dihydrate | 0.00025 | Micronutrient/vitamin |
| Potassium iodide | 0.00083 | Micronutrient/vitamin |
| Glycine | 0.002 | Micronutrient/vitamin |
| Myo-inositol | 0.1 | Micronutrient/vitamin |
| Nicotinic acid | 0.0005 | Micronutrient/vitamin |
| Pyridoxine HCl | 0.0005 | Micronutrient/vitamin |
| Thiamine HCl | 0.0001 | Micronutrient/vitamin |
| Zinc sulfate heptahydrate | 0.0086 | Zinc solution |
| Iron sulfate heptahydrate | 0.0278 | Iron sulfate solution |
| EDTA | 0.0296 | EDTA solution |

Macronutrient solution was prepared as 1x solution and adjusted to pH 5.8 with KOH. The solution was autoclaved for 20 min at 120 °C.

Micronutrient/vitamin solution was prepared as 500x stock solution and filter sterilized.

Iron sulfate, EDTA and zinc solution were prepared as 1000x stock solution. EDTA solution was adjusted to pH 8.0 with KOH. All solutions were filter sterilized. Iron sulfate solution was kept at -20 °C and defrosted prior to use.

**Table S3.** Additional oligos used in this study. OE, overexpression.

| Primer name | Sequence 5'→3' | Notes |
|---|---|---|
| SN01 | CAAATTCGCGACCGGTATGGCAAAGAACCTCAACTCC | Assembly primers for pEAQ-zdrp1.2 |
| SN02 | CCAGAGTTAAAGGCCTCGAGTTACGATTTGTAGCAATGGCAGTG | |
| SN03 | CAAATTCGCGACCGGTATGGCAAAGCTCATAGTTAACTTCTC | Assembly primers for pEAQ-zdrp2 |
| SN04 | CCAGAGTTAAAGGCCTCGAGTTAACAAGGTCCGTAGCAGTGAC | |
| SN05 | CAAATTCGCGACCGGTATGGCAAAGAACCTCAACACTG | Assembly primers for pEAQ-zdrp3 |
| SN06 | CCAGAGTTAAAGGCCTCGAGTCAAGACGTTCCGTAGCAATG | |
| SN07 | CAAATTCGCGACCGGTATGGCAAAGACTCAAAACTTTGTTTGC | Assembly primers for pEAQ-zdrp4 |
| SN08 | CCAGAGTTAAAGGCCTCGAGTTAGCGGCGACAATGACAATG | |
| SN09 | TTCTGCCCAAATTCGCGCACATGGCAAAGAACCTCAAC | Assembly primers for OE plasmid 1301Y-ZDRP1 |
| SN10 | CCAGAGTTAAAGGCACGTTTTTACGATTTGTAGCAATGGCAGTG | |
| SN11 | CCAGAGTTAAAGGCCTCGAGTCAACAATAGCAATGACAGTGTTTCTCACTTCC | Assembly primers for pEAQ-*Brapa*ZDRP and pEAQ-*Bnapus*ZDRP |
| SN12 | CAAATTCGCGACCGGTATGGCTAAGAACATCAACTCAGTCAGC | |
| SN13 | CTCAACACGTCCAGCACAAA | Confirmation of OE plasmid 1301Y-ZDRP1 |
| SN14 | CGGTGTGGTCTTGGGAAAAG | |

**Table S4.** Selected ZDRP sequences from *Brassica* spp. Q*, predicted pyroglutamic acid. Calc., calculated.

| Organism | Core sequence | Calc. mass | Accession number |
|---|---|---|---|
| *Arabidopsis lyrata subsp. lyrata* | ACFTFLGECGPEPFTGSNADCLACCVALYSSPPVCAGRVEGNPAHCHCYKS | 5281.2 | XP_002878305.1 |
| | ACFKFLGECGAVPFTGSNADCKSCCEGKFGSAAVCAGRVEAEGGVNHCHCYGTS | 5434.3 | XP_002891073.1 |
| *Camelina sativa* | Q*TCSTFLDECGPDPFLGTNADCYNCCKYTYGSPPACKGLVEGSDNHCHCYQNP | 5748.3 | XP_010470473.1 |
| | Q*TCSTFLDECGPAPFLGTNADCFKCCKYTYGSPPACKGLVEGSDKHCHCYQNP | 5716.4 | XP_010513408.1 |
| | Q*TCSTFLDECGPEPFLGANADCFNCCKYTYESPPACKGLVEGSDKHCHCYQNP | 5802.4 | XP_010424473.1 |
| | Q*TCSTFLDECGPDPFLGTNADCFNCCKYTYGSPPACKGLVEGSDKHCHCYQNP | 5746.4 | XP_010456819.1 |
| | APACFTFLDECGPDPFPGTNVDCTKCCRSTYKGQHVCKGVVEGSENHCHCYQRTKKE | 6322.8 | XP_010514830.1 |
| | Q*TCSTFLDECGPEPFLGTNADCFNCCKYTYGCPPACKGLVEGSDKHCHCYQNP | 5776.4 | XP_019085758.1 |
| | AEPACFTFLDECGPDPFPGTNVDCTKCCRSTYKGQHVCKGVVEGSENHCHCYQRTKKE | 6451.8 | XP_010503150.1 |
| | APGCFTFLEECGPAPFLGTNVDCSICCRRTYNGQKVCKGVVEGSDKHCHCYQRTKEE | 6312.8 | XP_019101084.1 |
| | Q*TCSTFDECGPDPFLGTNSDCFNCCKYTYGSPPACKGLVEGSDKHCHCYQNP | 5649.3 | XP_010470521.1 |
| | Q*TCSTFLDECGPAPFLGTNADCLKCCKYTYGSPPACKGLVEGSDKHCHCYQNP | 5682.5 | XP_019093272.1 |
| *Capsella rubella* | AEPRCFTFLDECGPDPFLGTNLDCTECCRSTYPGQKVCKGVVEGSEQHCHCYQRSKKE | 6527.9 | XP_006292967.1 |
| *Brassica oleracea var. oleracea* | Q*TFCFECGPEPFLGTNADCFNCCKTKYGSPPVVSGVVEGSEKHCHCYC | 5203.2 | XP_013595457.1 |
| | Q*TFCFECGPVPFLGTNADCFNCCKTKYGSPPVVSGVVEGREKHCHCYC | 5242.3 | XP_013593698.1 |
| | Q*TFCFECGPVPFLGTNADCFNRCKTKYGSPPVVSGFVEGSEKHCHCYC | 5274.3 | XP_013595402.1 |
| | Q*TFCFEGGPVPFLGTNADCFNCCKTKHGSPPVVSGVVEGSEKHCHCYC | 5101.2 | XP_013595413.1 |
| *Raphanus sativus* | Q*TFCFECGPVPFLGTNADCFNCCKTKYGSPPVVSGVVEGSEKHCHCYC | 5173.2 | XP_018438468.1 |
| | Q*TFCFECGPVPFLGTNADCFNCCKTKYGSPPVVSGVLEGSEKHCHCYC | 5187.2 | XP_056844535.1 |
| | Q*TFCFECGPVPFLGTNADCFNCCKTKYGSPPVVSGVVEGSEKHCHCYC | 5173.2 | XP_056844507.1 |
| *Brassica napus* | Q*TFCFECGPVPFLGTNADCFNCCTTKYGSPPVVSGVVEGSEKHCHCYC | 5146.2 | XP_013682305.1 |
| | Q*TFCFECGPVPFLGTNADCFNCCKTKYGSPPVVSGFVEGSEKHCHCYC | 5221.2 | XP_013682102.1 |
| | Q*TFCFECGPVPFLGTNADCFNCCKTKYGSPPVVSGVVEGSEKHCHCYC | 5173.2 | XP_013718688.2 |
| | Q*TFCLECGPVPFLGTNADCFNCCKTKYGSPPVVSGVVDGSETHCHCYC | 5098.2 | XP_013681410.1 |
| *Eutrema salsugineum* | DQRTNSRCLPRGCKNPVFSEECGPEPFTGSNNDCCHCCIAKYGRKSVCKGVVEGSAKHCHCYKESV | 7250.2 | XP_024004825.1 |
| | ADRRTNSRCLPRGCTNPVFSEECGPEPFTGSNNDCCHCCIAKYGKKSVCKGVVEGSAKHCHCYKESV | 7294.3 | XP_024005193.1 |
| | ADQRTNSRCLPRGCTNPVFSEECGPEPFTGSNNDCCHCCIAKYGRKSVCKGVVEGSAKHCHCYKESV | 7294.2 | XP_024005179.1 |

### Supplemental Figures

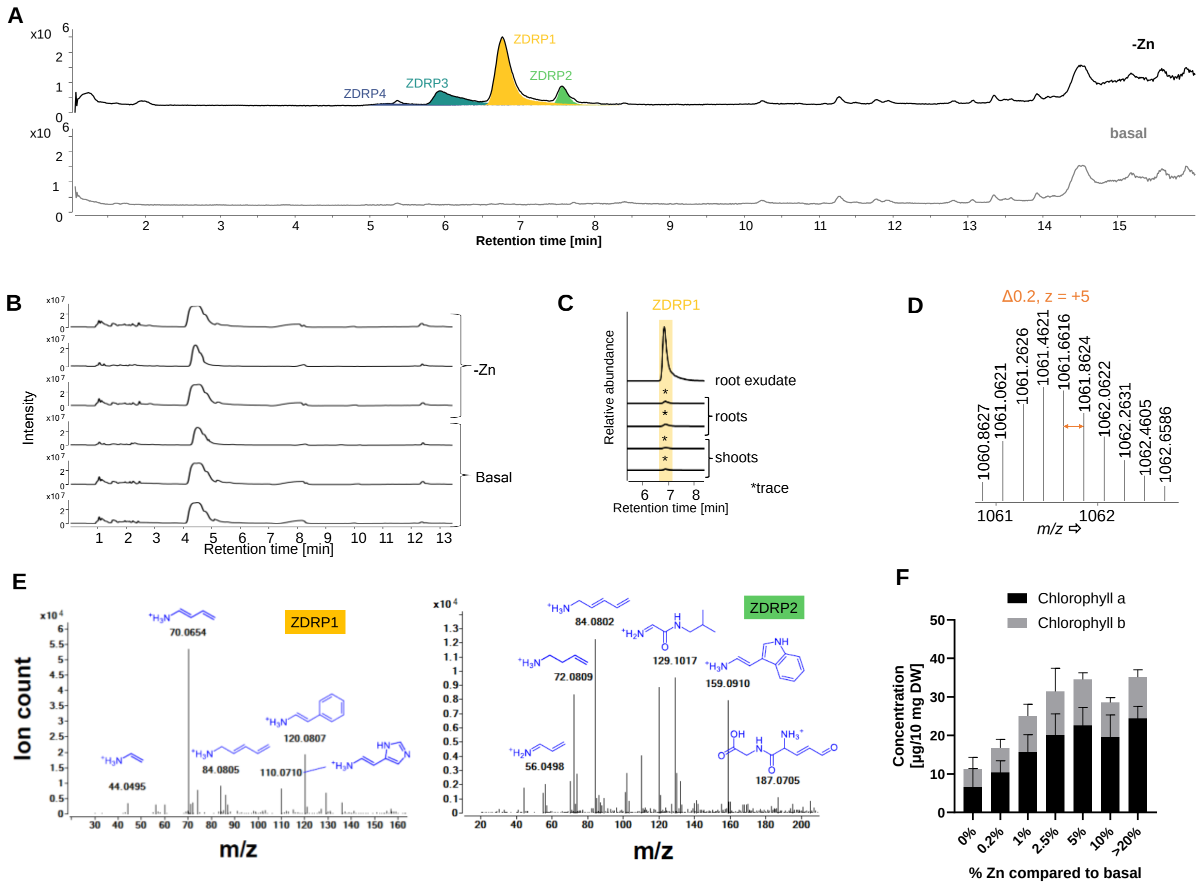


Figure S1. **ZDRPs are being produced in Zn-deficient soil.** A) Extended metabolic profiles of root exudates from *A. thaliana* grown under -Zn or basal conditions. RP-LC/MS in positive mode. Total ion chromatograms, TICs. B) Metabolic profiles of root exudates from *A. thaliana* grown under -Zn or basal conditions investigated by normal-phase LC/MS (HILIC) in positive mode. C) Trace amounts of ZDRP1 detected in shoot and root tissue of *A. thaliana* grown under -Zn vs. ZDRP1 level in root exudates. EIC, extracted ion chromatograms. D) Isotopic pattern of ZDRP1 as multi-charged mass signature (z=5) detected in soil extracts. E) Part of MS/MS fragmentation profile of ZDRP1 and ZDRP2. F) Chlorophyll content of Col-0 plants grown under various amounts of Zn. Tissue harvest after 28 days.

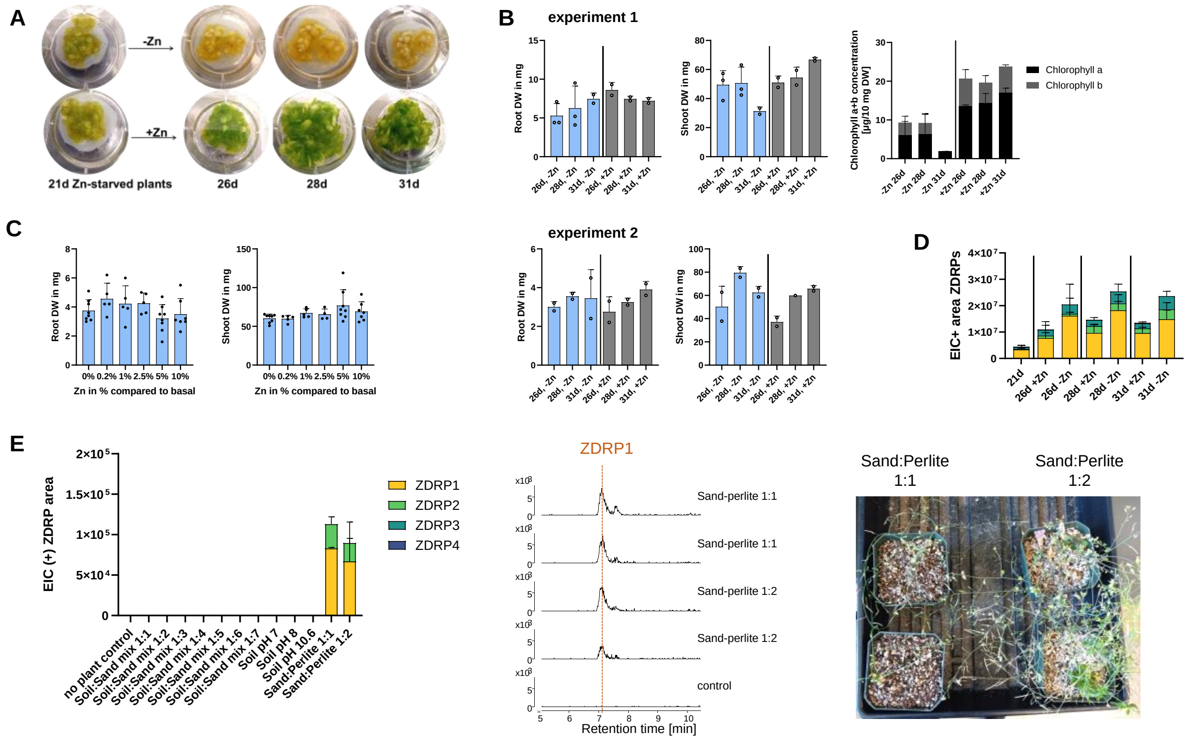


Figure S2. **Zn content influences root and shoot weights in plants.** A) Rescue of yellowing phenotype observed with Zn deficiency after resupply of basal zinc. B) Zinc re-supply leads to a rapid increase in shoot weights. Shown are chlorophyll content, root and shoot dry weights of *A. thaliana* var. Col-0 grown in hydroponics in two independent experiments. Harvest after 28 days. C) Zinc gradient reveals an increase in shoot dry weight with increasing zinc(II) concentration. Shown are root and shoot dry weights (DW) after 28 days grown in hydroponics. D) LC/MS analysis of ZDRP levels from plants re-supplied with Zn (+Zn) or continuously grown without Zn (-Zn). E) LC-MS analysis of ZDRPs in soil extracts produced by *A. thaliana*. Control corresponds to *Arabidopsis* plants grown on conventional soil with slow-release fertilizer. *A. thaliana* var. Col-0 plants grown in sand:perlite mixtures for 6 weeks.

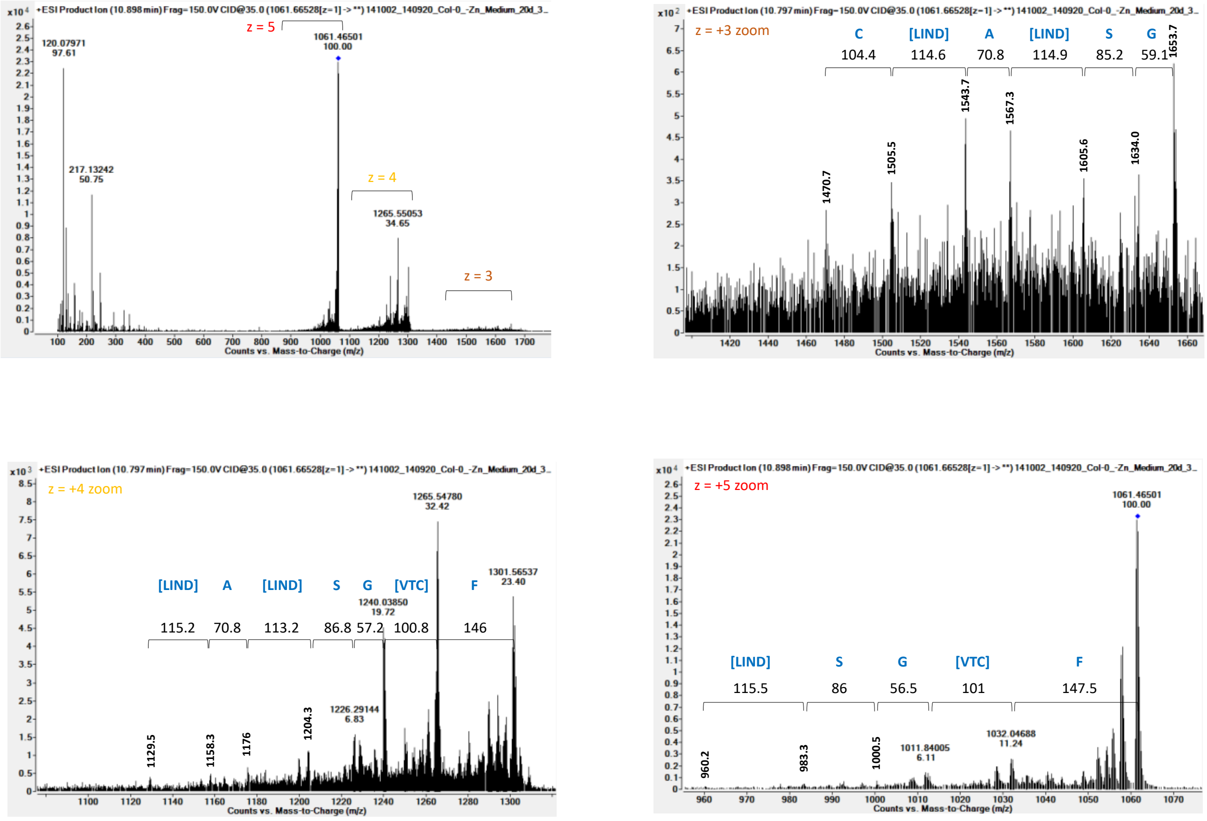


Figure S3. **MS/MS fragmentation pattern led to the biosynthetic origin of ZDRP1.** Fragmentation pattern of ZDRP1 in positive mode. Possible amino acid fragments are indicated in brackets.

**A**

| Gene name ⇨ / Transcriptomics study/ Proteomics study ⇩ | At3g59930, At5g33355 | At2g36255 | At1g34047.1 | At4g11393 |
|---|---|---|---|---|
| Chen *et al*., 2018 | ✓ | ✓ | ✓ | ✓ |
| van de Mortel *et al*., 2006 | ✓ | × | × | × |
| Zargar *et al*., 2015 | × | ✓ | ✓ | × |
| Inaba *et al*., 2015 | ✓ | ✓ | ✓ | × |
| Arsova *et al*., 2019 | ✓ | ✓ | ✓ | ✓ |
| Nakayama *et al*., 2020 | ✓ | ✓ | ✓ | ✓ |
| Nishida *et al*., 2017 | × | ✓ | ✓ | ✓ |

**B**

| **ZDRP1** | **ZDRP2** | **ZDRP3** | **ZDRP4** |
|---|---|---|---|
| Cys1-Cys6 | Cys1-Cys8 | Cys1-Cys6 | Cys1-Cys7 |
| Cys2-Cys4 | Cys2-Cys5 | Cys2-Cys4 | Cys2-Cys5 |
| Cys3-Cys7 | Cys3-Cys6 | Cys3-Cys7 | Cys3-Cys8 |
| Cys5-Cys8 | Cys4-Cys7 | Cys5-Cys8 | Cys4-Cys9 |
| | | | Cys6-Cys10 |

**C**

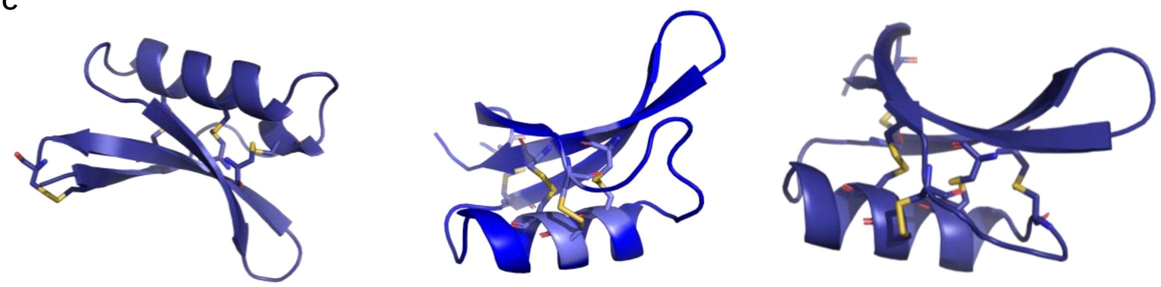

**ZDRP2 core peptide** **ZDRP3 core peptide** **ZDRP4 core peptide**

Figure S4. **Predictions of ZDRPs.** A) Transcriptomics and proteomics datasets mentioning *ZDRP1-4* upon Zn deficiency (16–22). B) Predicted cysteine disulfide patterns in ZDRP1-4 (by Alphafold). C) Structural prediction of ZDRP2-4 by AlphaFold. Core peptide in blue, disulfide bridges in yellow.

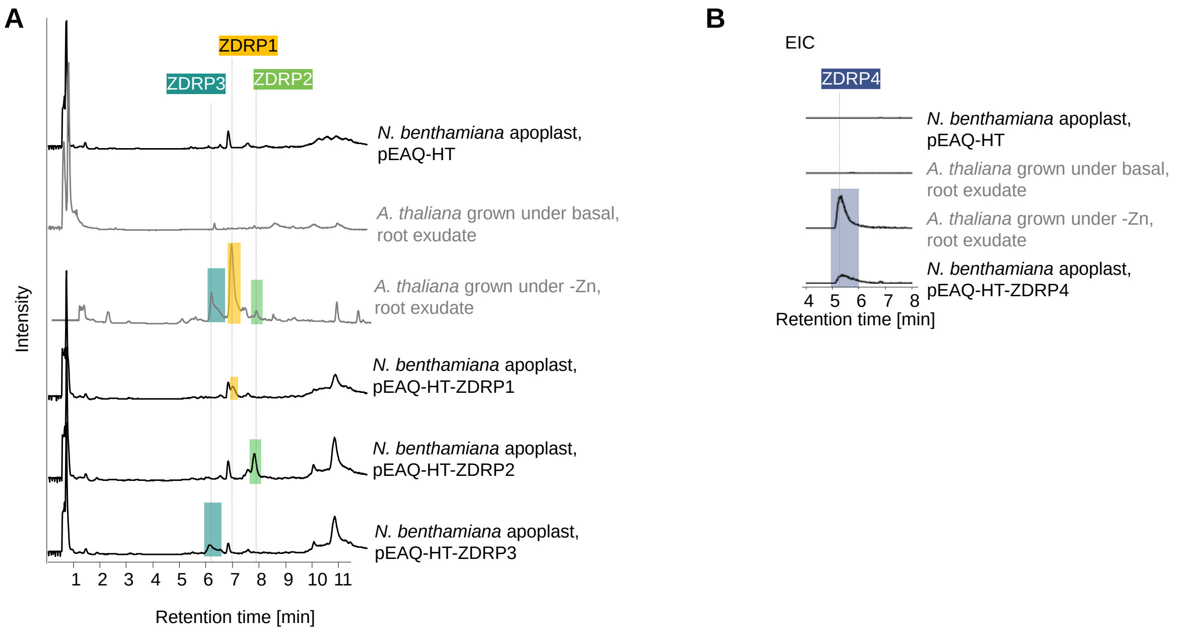


Figure S5. **Transient expression of ZDRP sequences reveal peptide production in *N. benthamiana***. A) Total ion chromatograms (TICs) of *A. thaliana* grown in hydroponics set-up (gray) or apoplasts of *N. benthamiana*. Heterologous expression of *ZDRP* genes in *N. benthamiana* lead to ZDRP production. *Agrobacterium*-mediated transformation was used to achieve peptide production in heterologous host (pEAQ-ZDRP1-3). B) *Agrobacterium*-mediated transformation was used to achieve peptide production in heterologous host (pEAQ-ZDRP4). ZDRP4 is only detectable as extracted ion chromatogram (EIC). All chromatograms in positive mode.

| Locus | Name | ZDRE sequence **RTGTCGACAY** | Position |
|---|---|---|---|
| At3g59930 | *ZDRP1.1* | GTGGCGACAT | -195 |
| | | GTGTCGCCAT | -507 |
| At5g33355 | *ZDRP1.2* | GTGGCGACAT | -192 |
| | | GTGTCGCCAT | -501 |
| At2g36255 | *ZDRP2* | ATGGCGACAT | -201 |
| At1g34047 | *ZDRP3* | GTGGCGACAT | -165 |
| | | GTGTCGACAT | -461 |
| At4g11393 | *ZDRP4* | ATGTCGACAT | -190 |
| | | ATGTTGACAT | -1014 |
| | | ATGTGGACAT | -1043 |

Figure S6. **Selected ZDRE promoter**. Zinc deficiency responsive element (ZDRE) in the promoter sequence of *ZDRP* genes. Highlighted in red are deviations from ZDRE motif.

A

| Gene | ZDRP | High impact variant numbers, Polymorph 1001 | Missense variant number, moderate, Polymorph 1001 | Mutation in signaling seq. (ePLANT) | Mutation in core seq. (ePLANT) |
|---|---|---|---|---|---|
| At3g59930 | **ZDRP1.1** | - | 57 | 50 | 7 |
| At5g33355 | **ZDRP1.2** | 5 | 5 | 5 | - |
| At2g36255 | **ZDRP2** | 2 | 1,466 | 351 | 1,119 |
| At1g34047 | **ZDRP3** | 28 | 352 | n.d. | n.d. |
| At4g11393 | **ZDRP4** | 1 | 2 | 484 | 20 |
| At1g10970 | **ZIP4** | - | 197 | n.d. | n.d. |
| At2g32270 | **ZIP3** | - | 565 | n.d. | n.d. |
| At1g05300 | **ZIP5** | 1 | 1,474 | n.d. | n.d. |
| At1g60960 | **IRT3** | 4 | 1,348 | n.d. | n.d. |
| At2g16770 | **bZIP23** | 11 | 302 | n.d. | n.d. |
| At4g35040 | **bZIP19** | - | 4 | n.d. | n.d. |
| At5g04950 | **NAS1** | 1 | 262 | n.d. | n.d. |
| At5g56080 | **NAS2** | - | 820 | n.d. | n.d. |
| At1g09240 | **NAS3** | - | 422 | n.d. | n.d. |
| At1g56430 | **NAS4** | 1 | 1,148 | n.d. | n.d. |

B

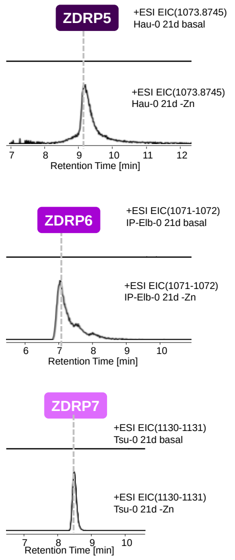


C

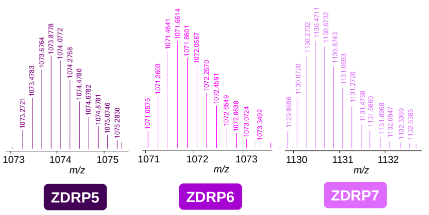


D

3x G→D; 1x C→Y; 3x V→L
ZDRP1.1 ACFTFLGECGPEPFTGSNADCLACCVALYKSPPVCAGRVEGVPAHCHCYKS

ZDRP1.2 ACFTFLGECGPEPFTGSNADCLACCVALYKSPPVCAGRVEGVPAHCHCYKS

1x C→S; 4x V→A; 1,097x R→K; 14x G→R
ZDRP2 Q*TCFKGEAQEGVCVKVDGSKLCDLLCRATNTTWFGACEVEDNETHCHCYGPC

ZDRP3 ACFKFLGECGAVPFPGTNADCTSCCVGNFGSAVCAGRVEVEGGVKHCHCYGTS n.d

1x T→P; 17x E→K; 1x K→I; 1x G→R
ZDRP4 KTCKLFRGECPVDPCEPEKCDECCKATFGKQICGKCEQESTELHCHCRR

E

| id | name | country | latitude | longitude | notes |
|---|---|---|---|---|---|
| 7411 | Wl-0 | GER | 47.9299 | 10.8134 | mutation in at1g34047, frameshift variant, p.Asn6fs/c.16_22delAACACTG |
| 6172 | TÅD | SWE | 62.8717 | 18.3436 | mutation in at1g34047, frameshift variant, p.Lys3fs/c.8delA |
| 9540 | IP-Elb-0 | ESP | 41.81 | 2.34 | mutation in at1g60960 (irt3), 2x frameshift variant, p.Thr40_Ser41fs/c.118_119insT, p.Ala39fs/c.116delC |
| 9774 | Alt-1 | GER | 48.59 | 9.22 | mutation in at2g16770 (bZIP23), frameshift variant, p.Val231_Asn232fs/c.692_693insT |
| 7077 | Co-1 | POR | 40.12 | -8.25 | mutation in at4g11393, stop gained, p.Gly37*/c.109G>T |
| 9590 | IP-Trs-0 | ESP | 43.37 | -5.49 | mutation in at5g33355, frameshift variant, p.Leu15_Val16fs/c.45_46insC |
| 9708 | Kardz-1 | BUL | 41.62 | 25.35 | mutation in at5g33355, frameshift variant, p.Leu17_Leu18fs/c.49_50insAA |
| 7164 | Hau-0 | DEN | 55.675 | 12.5686 | previous dataset/study Campos *et al.*, *J. Exp. Botany*, **2017**. |
| 7373 | Tsu-0 | JPN | 34.43 | 136.31 | previous dataset/study Campos *et al.*, *J. Exp. Botany*, **2017**. |
| 8354 | Per-1 | RUS | 58 | 56.3167 | previous dataset/study Campos *et al.*, *J. Exp. Botany*, **2017**. |

Figure S7. **ZDRP produced by *A. thaliana* ecotypes.** A) Occurrence frequency of mutations in Zn-deficiency upregulated genes. Data based of ePLANT and Polymorph 1001 (11). B) Additional ZDRPs being produced by *A. thaliana* ecotypes that are absent from Col-0 root extracts. Shown are extracted ion chromatograms of ZDRP5-7 from metabolic profiles of root exudates from plants grown in basal medium or under Zn deficiency (-Zn). C) Isotopic patterns of *z*=+5 ZDRP5-7 in positive mode. D) ePLANT prediction of mutations in core sequence of ZDRP proteins. ZDRP3 mutations were not determined since gene model 2.0 (At1g34047.2) is incorrectly assigned for this gene. E) Monitored ecotypes, their origin, and SNP of interest. Three ecotypes were chosen based on data from a previous study (23).

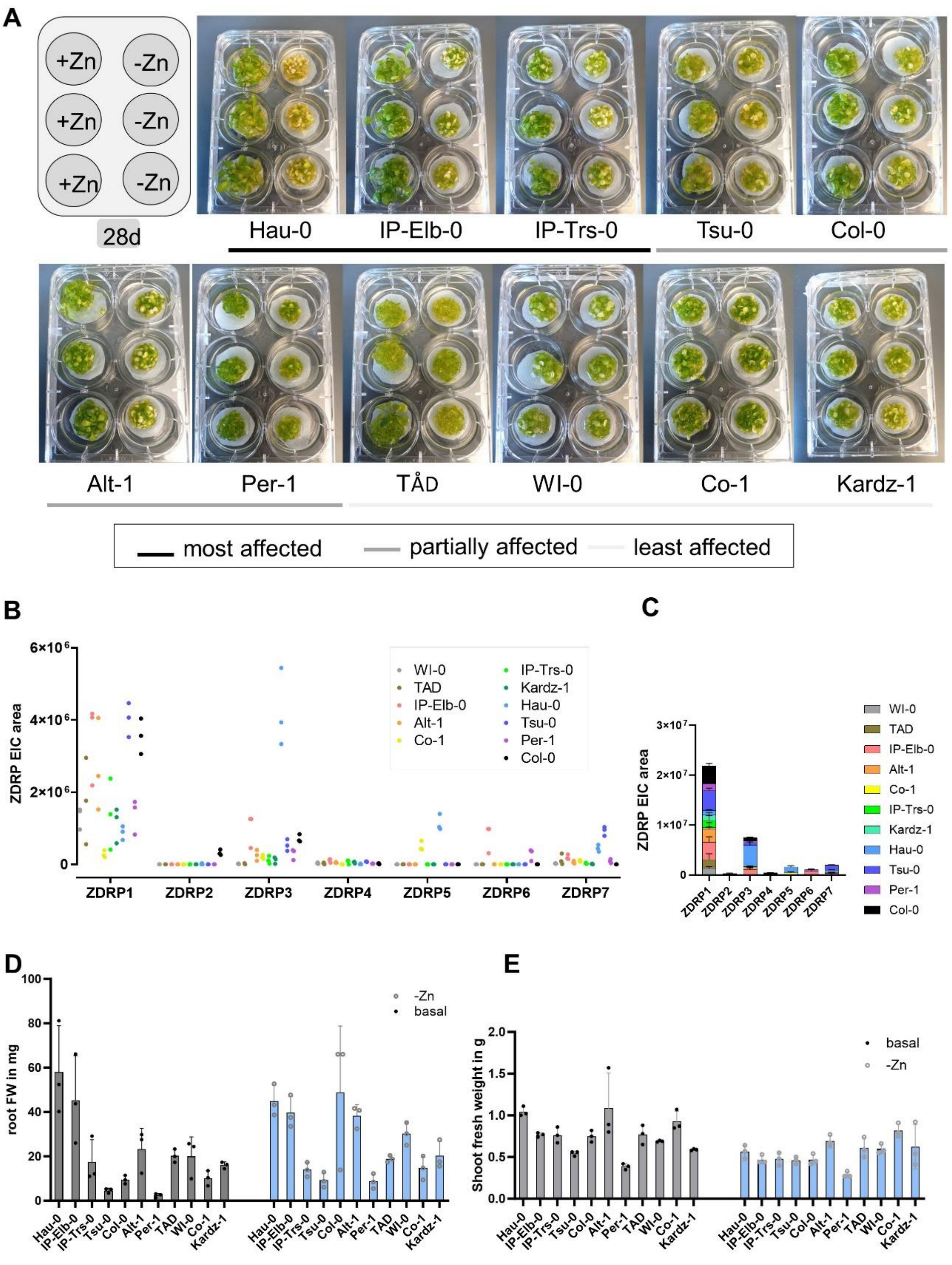


Figure S8. **ZDRP production from natural variants of *A. thaliana*.** A) Phenotype changes of ecotypes observed after 28 days. Plants affected by Zn deficiency show yellowing of leaves and stunted growth (most affected). Least affected plants showed no phenotype change when comparing basal with -Zn conditions. B) ZDRP levels in ecotype root exudate from plants grown under zinc deficiency. C) Cumulative distribution of ZDRP1-7 levels among ecotypes. D) Root fresh weight (FW) and E) shoot FW of ecotypes under basal and -Zn conditions.

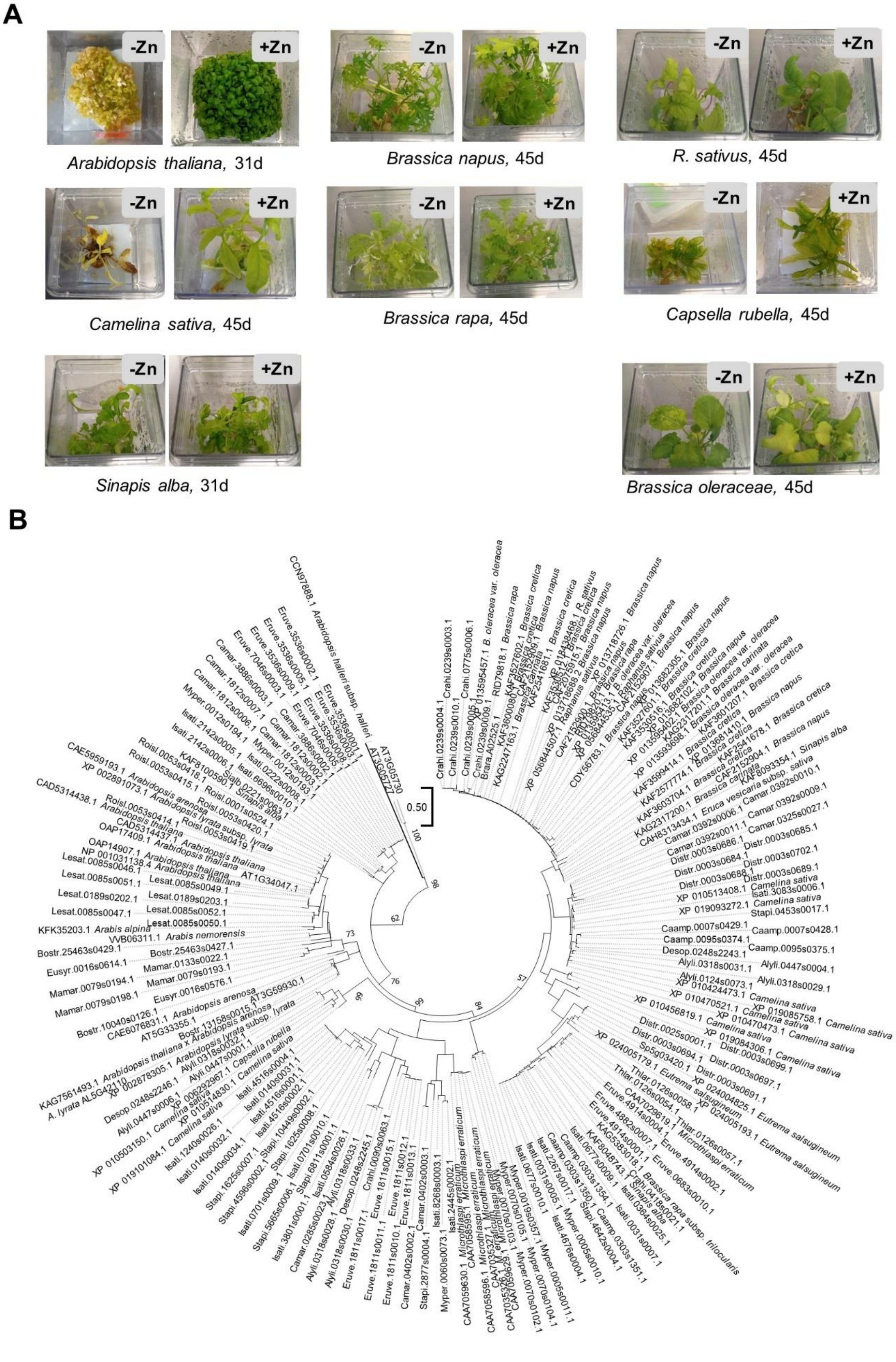


Figure S9. **Phylogenetic tree of ZDRP-like sequences.** A) Phenotype change of selected Brassicaceae plants grown under -Zn and basal conditions. B) Maximum likelihood tree of BLASTp results of amino acid sequence of proZDRP1.1. Bootstrapping 100, values under 50 were omitted. Sequences originated from Phytozome and NCBI database.

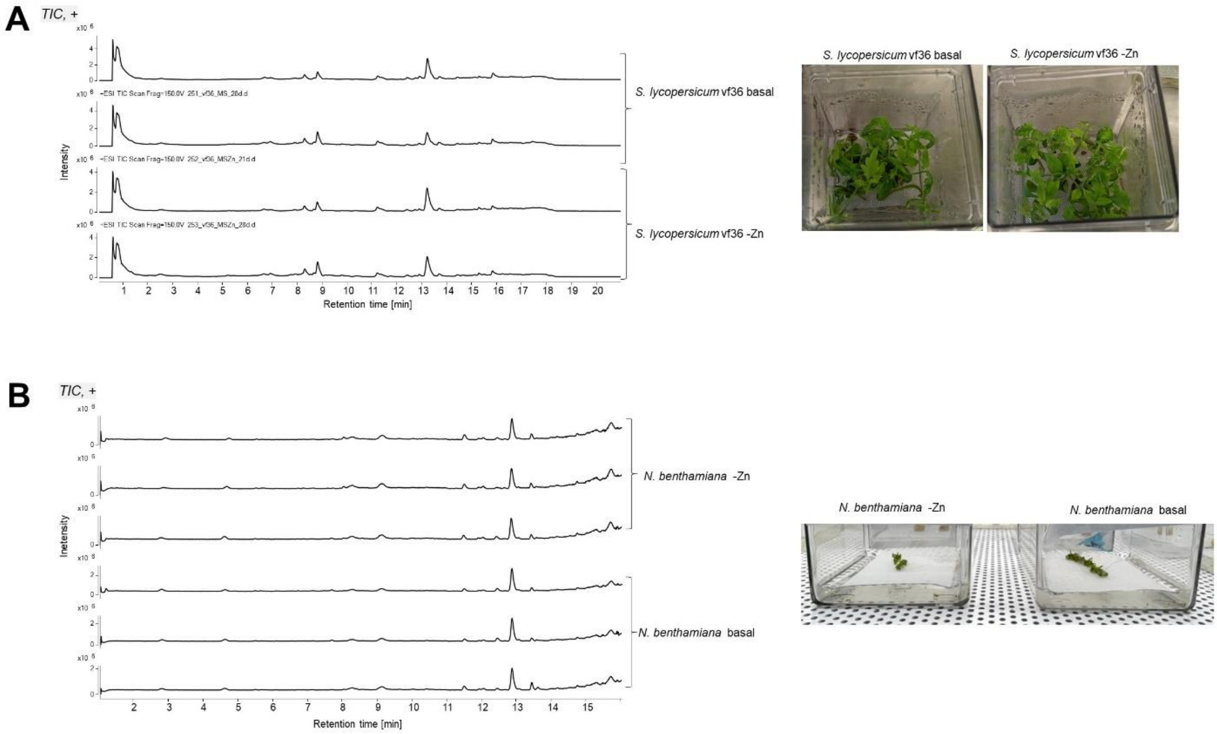


Figure S10. **Hydroponics set up with tomato (*S. lycopersicum*) and *N. benthamiana*.** A) Total ion chromatogram (TIC) of root exudates from *S. lycopersicum* grown under -Zn and basal conditions. Extracts taken after 21 days. Phenotype of tomato after 17 days. B) TIC of root exudates from *N. benthamiana* grown under -Zn and basal conditions. Phenotype of plants after 15 days.

**A**

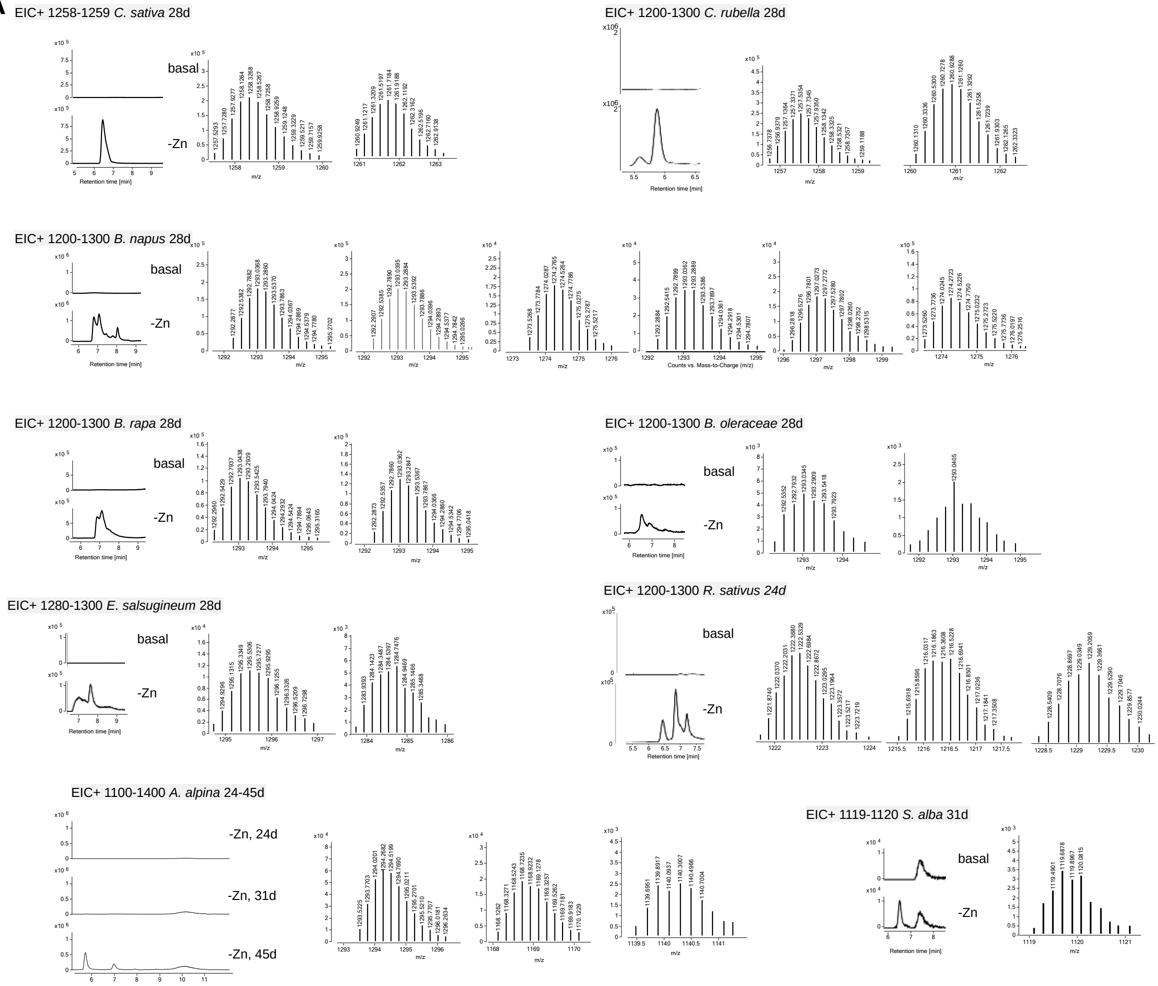


**B**

| Organism | Observed monoisotopic mass in positive mode | *z* |
|---|---|---|
| *A. alpina* | 1293.5225,* 1168.1262, 1139.5 | 4-5 |
| *B. napus* | 1292.2829,* 1292.2907, 1273.5268, 1292.2884, 1296.2818, 1273.5260 | 4 |
| *B. oleraceae* | 1291.77, 1292.276* | 4 |
| *B. rapa* | 1292.2960, 1292.2848* | 4 |
| *C. rubella* | 1256.7256,* 1260.1310 | 5 |
| *C. sativa* | 1257.5215,* 1260.9249 | 5 |
| *E. salsugineum* | 1294.9296, 1283.7400* | 5 |
| *R. sativus* | 1221.8716,* 1215.6918, 1228.5409 | 5 |
| *S. alba* | 1119.095* | 5 |

* most prominent ZDRP per root exudate

**C**

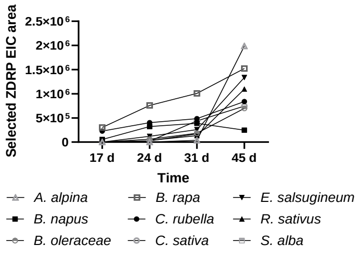


Putative *B. rapa* peptide:
Signal MAKNINSVSITVLLFVLLVVSTEILKSEA
Core Q*TFCFECGPVPFLGTNADCFNCCQTKYG
SPPVVSGVVEGSEKHCHCYC

**Figure S11. Extracted ion chromatograms revealing ZDRP-like production under Zinc deficiency in root exudates of plant-of-interest.** A) Metabolic profiles of root exudates from plants grown under basal or -Zn conditions are compared. ZDRP ion patterns (*z=4-5*) are highlighted. Positive mode peptide detection with LC/MS. B) Overview of detected peptides. C) Time-dependent production of most abundant ZDRPs in Brassicaceae cultures and putative identified *B. rapa* peptide. Most abundant ZDRP is marked in Figure S11B. Q*, predicted pyroglutamic acid.

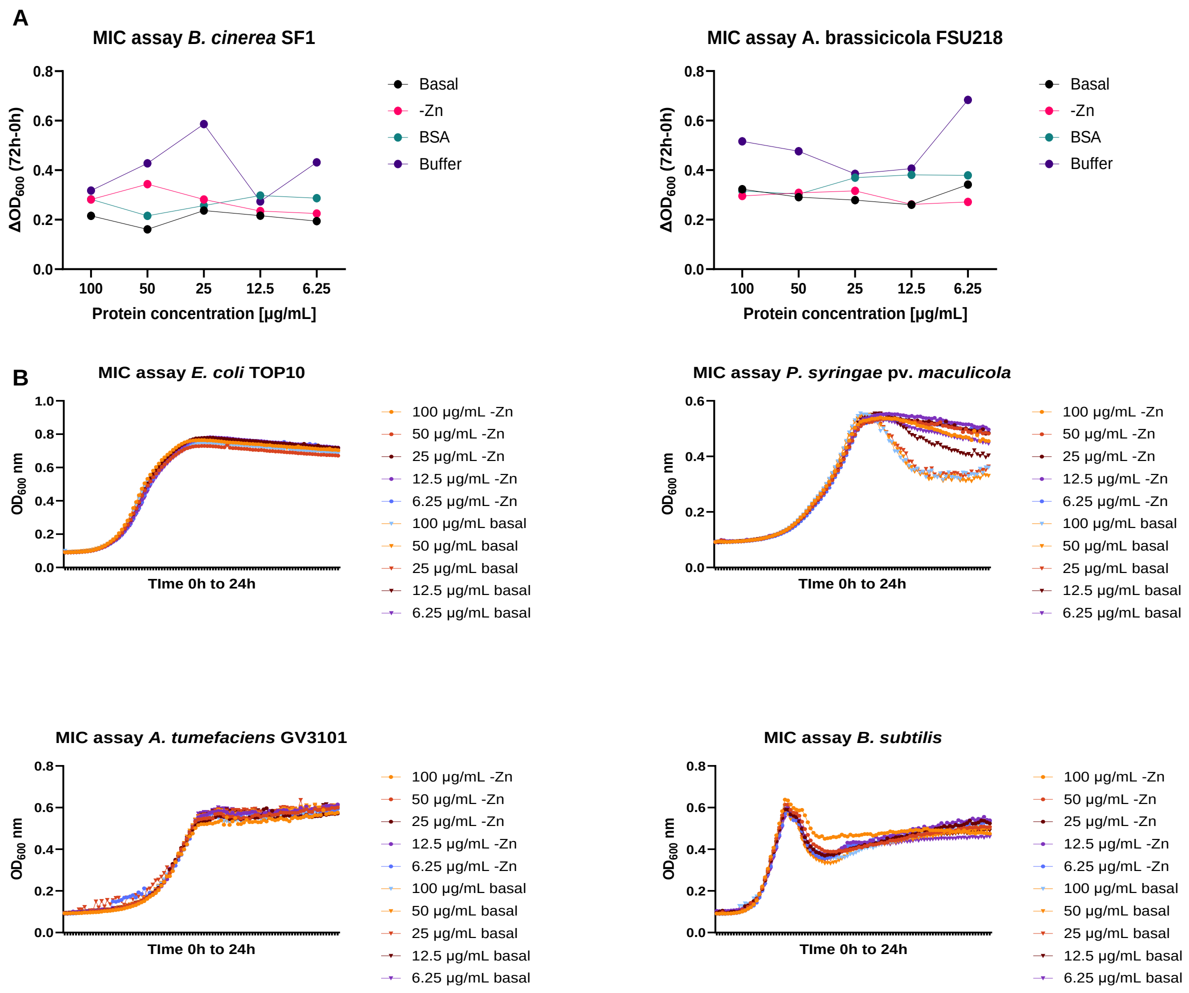


Figure S12. **Antimicrobial assays reveal ZDRPs are not antibacterial or antifungal (against the tested strains).** A) Growth curve of two fungal strains grown in the presence of increasing peptide amounts harvested from basal or -Zn root exudates. B) Growth curve of four bacterial strains grown in the presence of increasing peptide amounts harvested from basal or -Zn root exudates.

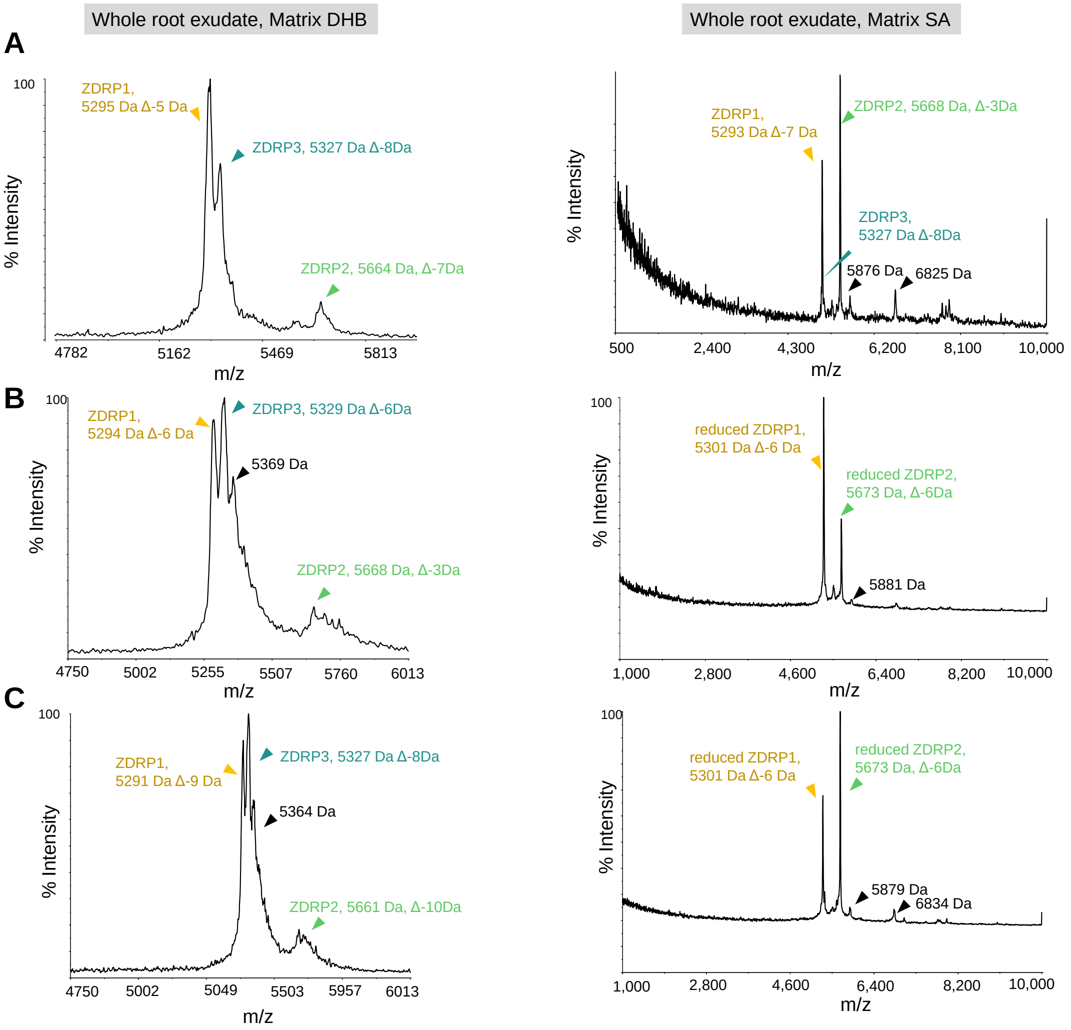


Figure S13. **MALDI-TOF/MS analysis of root exudate reveals reduced peptide form and no zinc binding.** A) Oxidized form of ZDRPs observed with MALDI-TOF in 2,5-dihydroxybenzoic acid (DHB) or sinapinic acid (SA) matrix. B) Reduced form of ZDRPs observed with MALDI-TOF in SA but not DHB matrix after TCEP treatment. C) Reduced form of ZDRPs observed with MALDI-TOF in DHB or SA matrix after TCEP and Zn treatment. Both matrices reveal ZDRPs to be the most prevalent peptide/protein species in -Zn root exudates. Δ, difference between theoretical and observed mass.

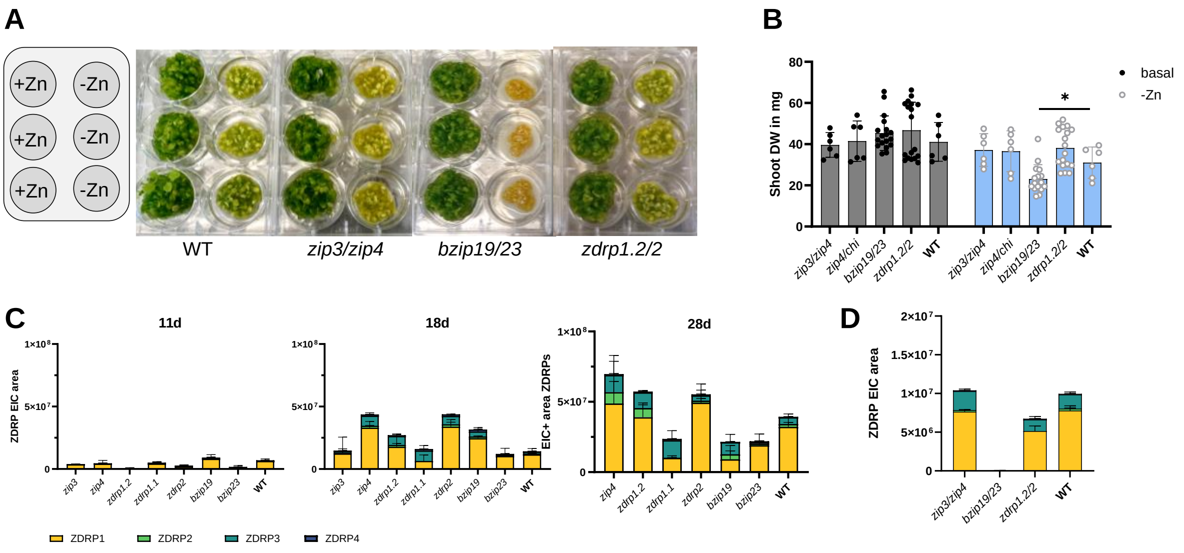


Figure S14. **Mutant experiments support ecological role in root development.** A) Phenotype change of T-DNA double mutants under basal vs -Zn conditions. B) Shoot dry weight (DW) of double mutant plants grown under basal or -Zn conditions. C) LC/MS analysis of single mutants grown under -Zn after 28 days. D) LC-MS analysis of double mutant exudates from plants grown under -Zn conditions. Experimental repeat.

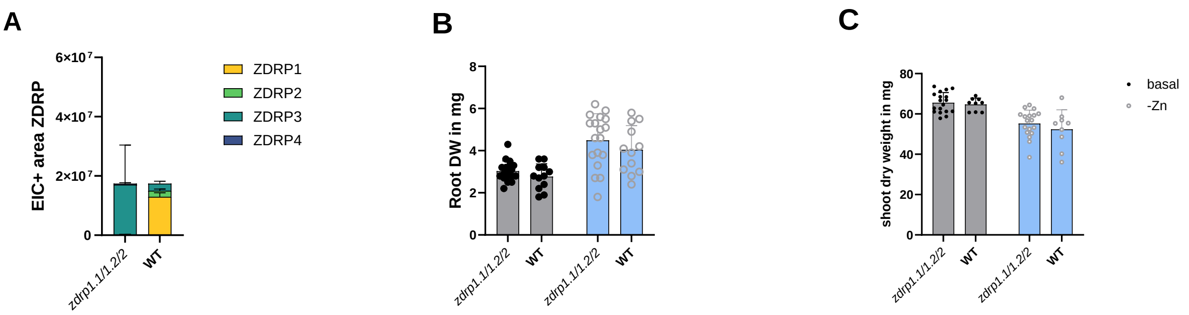


Figure S15. **ZDRP1/2-deficienct mutant shows complimentary effect.** A) LC/MS analysis of root exudates from plants grown under -Zn. ZDRP levels in *zdrp1.1/1.2/2* mutant compared to WT levels. B) Root DW of *zdrp1.1/1.2/2* mutant vs the WT after 28 days. C) Shoot DW of *zdrp1.1/1.2/2* mutant vs the WT after 28 days.

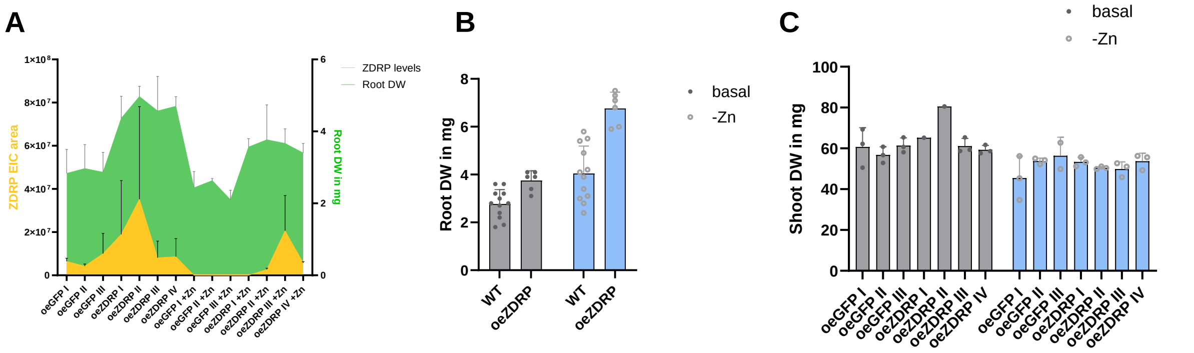


Figure S16. **Four independent ZDRP1-overexpression lines secrete ZDRP1 from roots to influence root development.** A) Correlation between ZDRP levels and root dry weight in overexpression strains oeGFP and oeZDRP. B) Experimental repeat: root DW of ZDRP1 overexpression lines and WT strains grown under -Zn or basal conditions after 28 days. C) Shoot dry weights of GFP and ZDRP overexpression strains grown in hydroponics under basal or -Zn conditions.

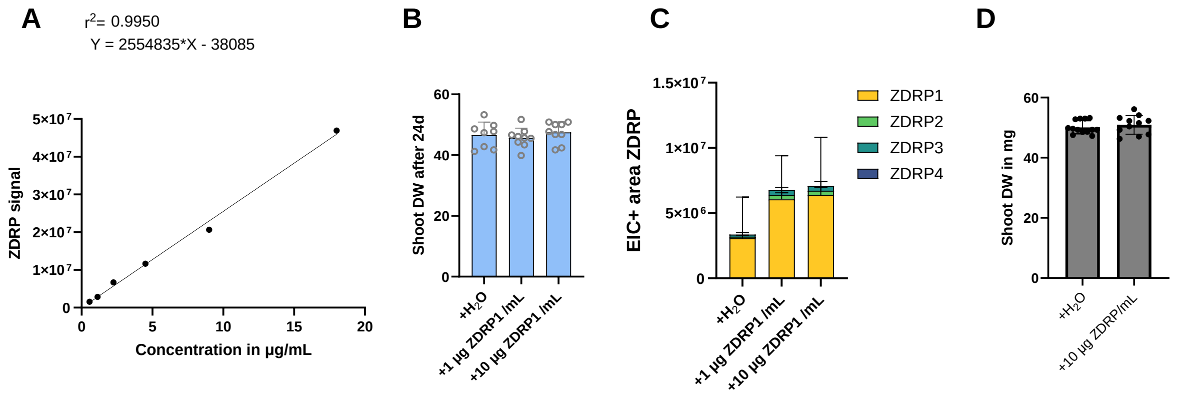


Figure S17. **Chemical complementation of ZDRP1 to WT strains enhances root weight.** A) Standard curve for ZDRPs. EIC signal in positive mode. B) Shoot dry weights of WT plants with or without ZDRP1 addition. Harvest after 24 days. Growth under -Zn conditions. C) Increased ZDRP levels in cultures supplemented with ZDRP1. D) Shoot dry weights (DW) of plants grown under basal conditions with or without addition of ZDRPs.

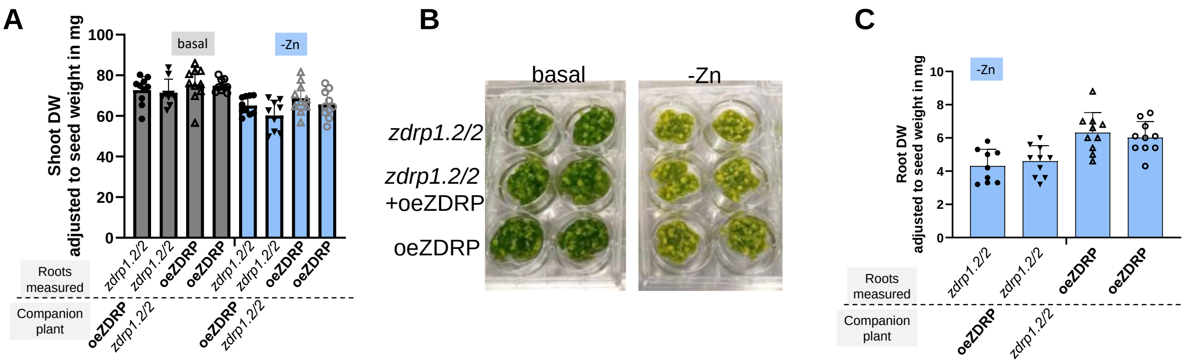


Figure S18. **Chemical complementation experiment.** A) Shoot DW from double mutant strain *zdrp1.2/2* and ZDRP overexpression strain oeZDRP grown alone or side-by-side. B) No phenotype changes observed for strains grown alone or in proximity to ZDRP overexpressing strain. C) Root DW from double mutant strain *zdrp1.2/2* and ZDRP overexpression strain oeZDRP grown alone or side-by-side under -Zn conditions.
